# Bayesian State-Space Model for Joint Inference of Oscillatory Dynamics and Point-Process Coupling

**DOI:** 10.64898/2026.06.15.732402

**Authors:** Bowen Zheng, Scott L. Brincat, Jacob A. Donoghue, Earl K. Miller, Emery N. Brown

## Abstract

Under a range of behavioral and physiological conditions, spike times and local field potential (LFP) oscillations exhibit phase coupling within specific frequency bands. Classical measures such as spike–field coherence (SFC) and the phase-locking value (PLV) quantify this coupling but estimate the LFP spectrum independently of spike timing. We introduce Joint SSMT, a Bayesian state-space framework that jointly infers LFP spectrograms and spike–field coupling strength. The model treats narrowband LFP activity as a latent process evolving in continuous time, with spike trains linked to the complex spectral state through a Bernoulli–logistic model. In simulations, Joint SSMT accurately recovers coupling strength, denoises the spectrogram, and uses spike timing to resolve fine temporal structure in the LFP. Applied to propofol anesthesia data, the model identifies coupling at a specific slow-oscillation frequency where SFC and PLV report only broad low-frequency coupling. We extend Joint SSMT to trial-structured experiments and apply it to primate recordings during an associative learning task, revealing frequency-specific coupling in hippocampus and prefrontal cortex. We also derive closed-form expressions for SFC and PLV as functions of the generative model parameters. Across simulations and two primate datasets, Joint SSMT provides more frequency-specific coupling estimates with principled uncertainty quantification than classical PLV and SFC.

## 1 Background and motivation

Across diverse brain areas and behavioral conditions, neural activity exhibits phase coupling between spike timing and LFP oscillations. In macaque visual cortex, attention to a stimulus enhances gamma-band synchronization between spikes and the local field potential Fries et al. [2001]. This coupling predicts the speed of behavioral responses on a trial-by-trial basis Womelsdorf et al. [2006]. In macaque prefrontal cortex during working memory, gamma bursts accompany the encoding and retrieval of items held in working memory, appearing selectively at recording sites where spiking carries stimulus information Lundqvist et al. [2016]. In the rat hippocampus, place cells fire at progressively earlier phases of the theta rhythm as the animal traverses a place field O’Keefe and Recce [1993]. Similar phase relationships have been documented in rat entorhinal cortex Hafting et al. [2008], medial prefrontal cortex Jones and Wilson [2005], and human medial temporal lobe during navigation Qasim et al. [2021]. Global brain state also shapes spike-field coupling. In macaques, propofol anesthesia entrains spiking to slow oscillations and disrupts higher-frequency coherence between cortical areas Bastos et al. [2021]. Together, these findings show that spike–field coupling is observed across species, brain areas, and behavioral states.

Several approaches have been developed to quantify the coupling between spike times and local field potential (LFP) oscillations. Spike-field coherence computes the spectral coherence between the spike train (treated as a point process) and the LFP signal Jarvis and Mitra [2001], Pesaran et al. [2002]. Spike-triggered average methods instead align LFP segments to each spike time, average across spikes, and quantify the oscillatory content of this average relative to the baseline LFP power Fries et al. [2001]. Phase-locking methods extract the instantaneous LFP phase at each spike time and summarize phase concentration using circular statistics. Bias-reduced metrics such as the pairwise phase consistency address the dependence of traditional measures on spike count Vinck et al. [2010, 2012].

These coupling measures share a common structure. They first estimate spectral quantities from the LFP, such as instantaneous phase or Fourier coefficients, and then relate spike timing to those estimates. The spectral quantities are treated as fixed, known values. Uncertainty in the spectrogram does not propagate into uncertainty in the coupling estimate. When the LFP is noisy or nonstationary, spectral estimates may be unreliable, and the resulting coupling statistics inherit that uncertainty in ways that are difficult to quantify.

Spectrogram estimation has its own extensive methodology for handling uncertainty. Multitaper analysis reduces the variance of spectral estimates by averaging over multiple orthogonal tapers Thomson [1982]. More recently, Kim et al. [2018] embedded multitaper analysis in a state-space framework. In this approach, Fourier coefficients at each frequency are modeled as noisy observations of a latent spectral state evolving over time. This state-space multitaper (SSMT) method yields smoothed spectrograms with confidence intervals.

The problems of spectral estimation and coupling inference have thus far been addressed separately. Point-process methods can infer oscillatory modulation from spike trains alone Arai and Kass [2017], but do not incorporate LFP measurements. Yet when spikes and LFP are coupled, each signal carries information about the other. Spike time informs the probable phase and amplitude of the LFP oscillation at that moment. A joint model could use spike timing to sharpen spectral estimates while propagating spectral uncertainty into the coupling estimate itself.

We develop **Joint SSMT**, a Bayesian model that jointly estimates the LFP spectrogram and spike-field coupling. We refer to the full spike–LFP model obtained by adding the Bernoulli spike likelihood with Pólya–Gamma augmentation as Joint SSMT. A latent spectral state—the complex Fourier coefficient at each frequency—evolves in continuous time as an Ornstein–Uhlenbeck process. Multitaper LFP measurements provide noisy linear observations of this state at coarse temporal intervals. A Bernoulli-logistic model links spike probability to the same latent state, evaluated at sufficiently fine temporal resolution such that multi-spike bins are rare. Because both data streams observe a shared latent trajectory, spike timing refines the spectrogram and spectral uncertainty propagates into the coupling estimate. Pólya–Gamma augmentation renders the spike likelihood conditionally Gaussian. Conditional on the auxiliary variables, a Kalman smoother fuses LFP and spike pseudo-observations. The model returns posterior distributions over both the spectrogram and the coupling parameters.

The paper is organized as follows. Section 2 presents the generative model, inference algorithm, and extension to trial-structured data. Section 3 evaluates the method on simulated data and neural recordings during propofol anesthesia. Section 4 tests the inference framework on simulated data with trial structures and applies it to neural recordings during an associative learning task. Section 5 derives closed-form connections between model parameters and other coupling measures.

## 2 Theory

### 2.1 Generative model

Multitaper spectral analysis reduces the variance of spectral estimates by averaging over multiple orthogonal tapers [Thomson, 1982]. Given a discrete-time signal sampled at interval Δ, the method applies *M* orthonormal Slepian tapers 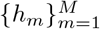 to a segment of *J* samples, computes the discrete Fourier transform of each tapered segment, and averages the resulting periodograms. Let *W* ∈ ℂ^*J*×*J*^ denote the unitary DFT matrix with entries (*W*)_*j*ℓ_ = *J*^−1*/*2^ exp[−2*πi*(*j* − 1)(ℓ − 1)*/J*], and let *ω*_*j*_ = 2*π*(*j* − 1)*/*(*J*Δ) denote the discrete angular frequencies for *j* = 1, …, *J*_*f*_, where *J*_*f*_ is the number of frequency bins of interest. For a tapered segment 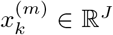 in time block *k*, the multitaper Fourier coefficient at frequency *ω*_*j*_ is 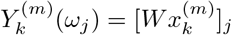.

Kim et al. [2018] embedded this spectral estimation procedure in a state-space framework. They modeled the complex Fourier coefficient at each frequency as a latent state that evolves over successive time blocks according to a random walk. For taper m, frequency *ω*_*j*_, and block index *k* ∈ ℕ, the model is

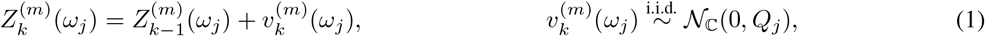

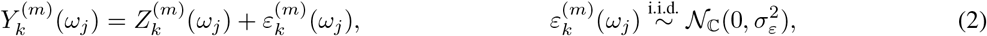

where *N*_ℂ_ (*µ, q*) denotes a circular complex normal distribution with mean *µ* and variance *q* = E[ *X* − *µ* ^2^]. Because tapers and frequency bins are conditionally independent, the model decomposes into *M* × *J*_*f*_ parallel one-dimensional complex Kalman filters. The EM algorithm recovers the process variances {*Q*_*j*_} and observation variance 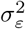 from data, yielding a smoothed spectrogram with pointwise confidence intervals and no hand-tuned smoothing parameters.

The state-space multitaper model operates on a block-by-block timescale. Resolving a narrowband spectral component at frequency *ω*_*j*_ requires a window spanning at least several oscillation cycles. In practice, the block width Δ_*b*_ = *J*Δ is often 100 ms or more. Within each block, the latent coefficient 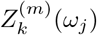 is treated as constant. Spikes, however, are typically binned at Δ_spk_ ≈ 5–10 ms and exhibit structure within a single spectral window. Coupling the spike train to the blockwise latent state would require either aggregating spikes within each window, which erases fine temporal structure, or shortening the window to match the spike timescale, which sacrifices frequency resolution. Neither compromise is satisfactory.

We address this mismatch by formulating the latent spectral state as a continuous-time stochastic process. For each taper *m* and frequency *ω*_*j*_, the complex Fourier coefficient evolves according to an Ornstein–Uhlenbeck diffusion,

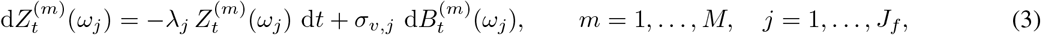

where 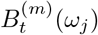 is a circular complex Wiener process. The parameter *λ*_*j*_ ≥ 0 governs the rate of mean reversion, where *τ*_*j*_ = 1*/λ*_*j*_ is the correlation time of the process. Setting *λ*_*j*_ = 0 recovers a complex Brownian motion. Discretizing this Brownian motion at the block spacing Δ_*b*_ yields exactly the random-walk dynamics in (1). For *λ*_*j*_ > 0, the process is stationary with marginal variance 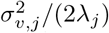. Because the diffusion is defined for all *t* ≥ 0, the latent state 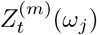 can be evaluated at any time—not only at block centers but also at the finer temporal resolution required for spike coupling.

At block centers *t*_*k*_ = *k*Δ_*b*_, multitaper Fourier coefficients provide Gaussian observations of the latent state,

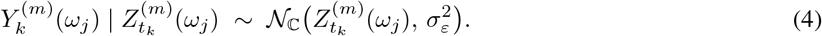

Independence across tapers and frequencies is preserved, so conditional on the spike data (discussed below), inference for the LFP-only model reduces to *M* × *J*_*f*_ parallel complex Kalman filters on the block grid. Without spike train observations, this model is a continuous-time reformulation of the state-space multitaper method described in Kim et al. [2018]. We refer to it as the continuous-time state-space multitaper method (CT-SSMT). Throughout the text, we use *CT-SSMT (LFP-only)* and *Joint SSMT* to distinguish the LFP-only model from the model that jointly incorporates both LFP and spike observations.

Spikes are recorded at a finer temporal resolution. We partition time into bins of width Δ_spk_ ≪ Δ_*b*_, with bin centers 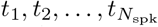. Each bin produces an indicator *S*_*n*_ ∈ {0, 1} denoting whether a spike occurred. To couple spikes with the latent spectral state, we first average the per-taper latent coefficients at each frequency and then rotate by the carrier phase to obtain the time-domain representation:

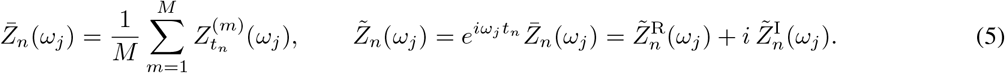

The quantity 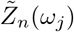 remains a linear function of the latent state because the phase factor 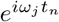 is deterministic. Its argument 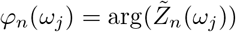 is the instantaneous LFP phase at frequency *ω*_*j*_, and its modulus 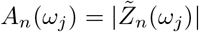 is the instantaneous amplitude.

We model spike probability as a function of instantaneous amplitude and phase across all frequency bands through a Bernoulli–logistic link. Conditional on the baseband latent states and spike history, the spike indicator follows

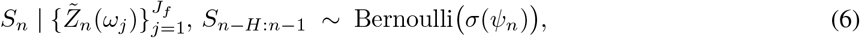

where *σ*(*u*) = 1*/*(1 + *e*^−*u*^) is the logistic function and the linear predictor is

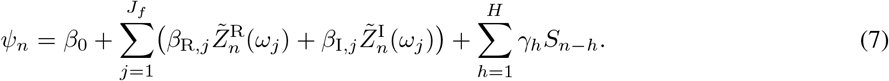

The parameters *β*_R,*j*_ and *β*_I,*j*_ capture phase-dependent modulation at frequency *ω*_*j*_. The spike history terms 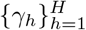 account for refractoriness and bursting. Writing *β*_C,*j*_ = *β*_R,*j*_ +*iβ*_I,*j*_ and 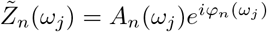 the linear predictor becomes

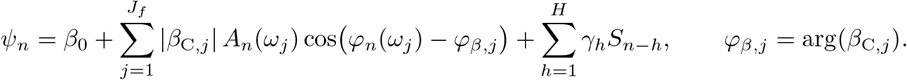

The modulus |*β*_C,*j*_| governs the depth of phase modulation at frequency *ω*_*j*_, *φ*_*β,j*_ is the preferred firing phase at that frequency, and *β*_0_ sets the baseline log-odds of spiking when all oscillation amplitudes are zero.

The logistic likelihood in (6) is not conjugate to the Gaussian prior on 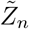 or to a Gaussian prior on the regression coefficients, precluding direct application of Kalman filtering. The next section introduces a data-augmentation strategy that restores conjugacy and enables efficient joint inference. Figure 1 summarizes the resulting inference pipeline.

**Table 1:** Summary of key notation. A complete notation reference appears in Appendix A.

| Symbol | Meaning |
| --- | --- |
| $t; t_n; t_k$ | Continuous time, spike-bin center, LFP block center |
| $\Delta_{\text{spk}}; \Delta_b$ | Spike-bin width, LFP block width |
| $n, k, m, j$ | Spike-bin, block, taper, frequency indices |
| $Z_t^{(m)}(\omega_j)$ | Complex latent OU state |
| $\tilde{Z}_n = e^{i\omega_j t_n} \bar{Z}_n$ | Phase-rotated baseband coefficient |
| $Y_k^{(m)}(\omega_j)$ | Multitaper LFP observation |
| $S_n \in \{0, 1\}$ | Spike indicator |
| $\lambda_j, \sigma_{v,j}^2$ | OU decay rate and diffusion coefficient |
| $\Phi_j, F_j$ | OU transition factor at block / spike-bin resolution |
| $\sigma_\varepsilon^2$ | LFP observation noise variance |
| $\beta_{C,j} = \beta_{R,j} + i\beta_{I,j}$ | Complex coupling coefficient |
| $\psi_n$ | Linear predictor (log-odds of spiking) |
| $\xi_n$ | Pólya–Gamma auxiliary variable |
| $\mathbf{D}$ | Design matrix for coupling regression |
| $X, \delta_r$ | Shared trajectory, trial-specific deviation |

**Figure 1:**
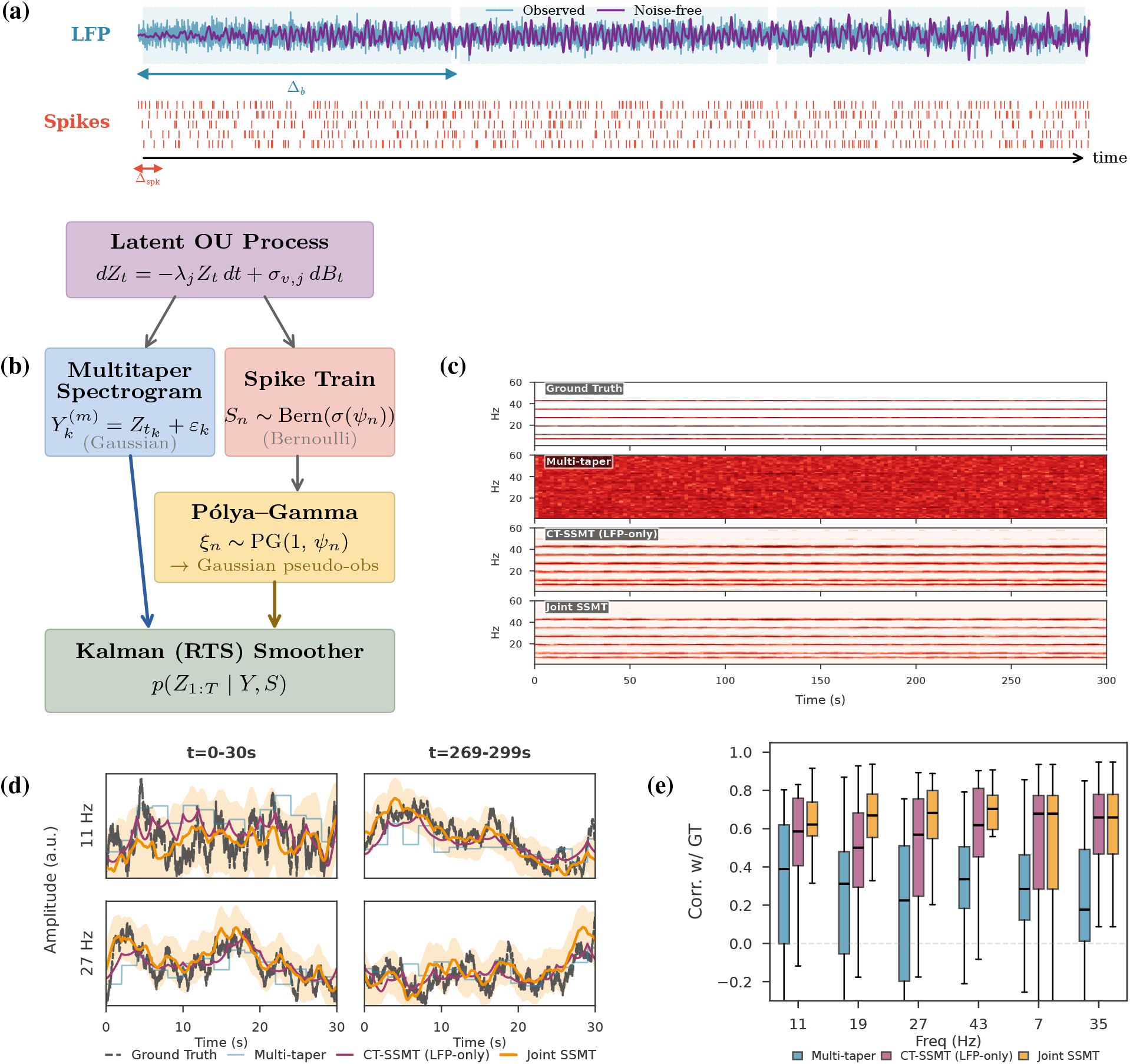
Joint SSMT framework for spike–field coupling. **(a)** Observed data: noisy LFP (blue) with underlying clean signal (purple) segmented into non-overlapping multitaper windows (Δ_*b*_), and multi-unit spike trains at fine temporal resolution (Δ_spk_). **(b)** Inference schematic: the latent spectral amplitude evolves as an Ornstein–Uhlenbeck process, generating Gaussian multitaper LFP observations and Bernoulli spike observations. Pólya–Gamma augmentation converts the spike likelihood to Gaussian pseudo-observations, enabling Kalman–RTS smoothing to compute *p*(*Z*_1:*T*_ |*Y, S*). **(c)** Spectrogram comparison across methods: ground truth (top), multitaper (second), CT-SSMT (LFP-only) (third), and Joint SSMT (bottom). **(d)** Time-resolved amplitude estimates at signal frequencies: Joint SSMT (orange) with 95% credible interval tracks ground truth (dashed black) more closely than multitaper (blue) or CT-SSMT (LFP-only) (magenta). **(e)** Correlation with ground truth power across 20-second windows, showing improved tracking by Joint SSMT.

### 2.2 Pólya–Gamma augmentation

The generative model comprises Gaussian observations (multitaper LFP coefficients) and Bernoulli observations (spike indicators), both linked to a shared latent spectral trajectory. The Gaussian component admits standard Kalman filtering, but the logistic link function in the spike model breaks conjugacy. The posterior over the latent state is no longer Gaussian. Pólya–Gamma augmentation resolves this problem by introducing auxiliary variables that render the Bernoulli likelihood conditionally Gaussian [Polson et al., 2013, Pillow and Scott, 2012]. We apply it here to couple spike observations with a latent spectral state that is jointly observed through multitaper LFP measurements.

We first describe inference when only LFP data are available. The Ornstein–Uhlenbeck diffusion (3) admits an exact discretization: over an interval of length Δ, the transition density is

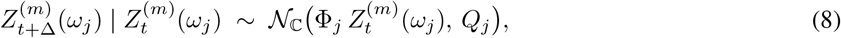

where 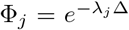 and 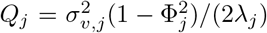 for *λ*_*j*_ > 0. At block centers *t*_*k*_ = *k*Δ_*b*_, multitaper coefficients provide Gaussian observations (4). Because tapers and frequencies are conditionally independent, inference reduces to *M* × *J*_*f*_ parallel one-dimensional complex Kalman filters. Rauch–Tung–Striebel smoothing yields posterior means and variances at each block center. The EM algorithm updates the parameters 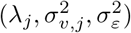 from smoothed sufficient statistics (Appendix B.1).

Incorporating spikes requires evaluating the latent state at the finer resolution Δ_spk_ ≪ Δ_*b*_. The OU dynamics can be discretized at this resolution, treating every spike-bin center *t*_*n*_ as a state time. The Bernoulli likelihood (6) is not conjugate to the Gaussian prior on 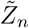 because 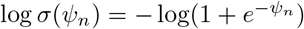 is not quadratic in *ψ*_*n*_. Pólya–Gamma augmentation restores conjugacy by representing the logistic function as a Gaussian scale mixture [Polson et al., 2013]. The Pólya–Gamma distribution PG(*b, c*) with *b* > 0 and *c* ∈ ℝ satisfies an integral identity for logistic functions: for any *b* > 0, *a* ∈ ℝ, and *ψ* ∈ ℝ,

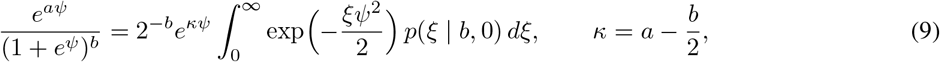

where *p*(*ξ* | *b*, 0) denotes the density of *ξ* ~ PG(*b*, 0). The right-hand side is a Gaussian kernel in *ψ* with precision *ξ*, averaged over a Pólya–Gamma mixing distribution. This identity converts a ratio of exponentials into a mixture of Gaussians, which is the mechanism that restores conjugacy.

The Bernoulli likelihood for spike bin *n* can be written as 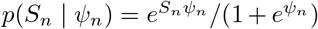, which matches the left-hand side of (9) with *a* = *S*_*n*_ and *b* = 1, giving 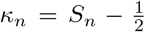. We introduce one auxiliary variable *ξ*_*n*_ per spike bin.

Conditional on *ξ*_*n*_, completing the square yields the augmented likelihood

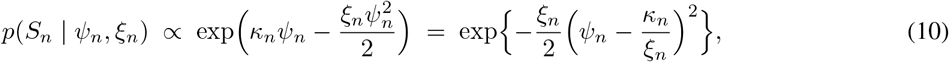

which is the kernel of a Gaussian in *ψ*_*n*_ with mean *κ*_*n*_*/ξ*_*n*_ and variance 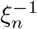. Equivalently, we may write

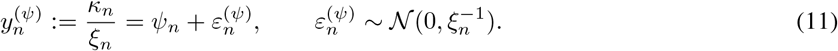

Each spike bin now contributes a Gaussian pseudo-observation of the linear predictor *ψ*_*n*_ with precision *ξ*_*n*_. The conditional distribution of the auxiliary variable is *ξ*_*n*_ | *ψ*_*n*_, *S*_*n*_ ~ PG(1, |*ψ*_*n*_|). Efficient samplers are available [Polson et al., 2013, Windle et al., 2014].

### 2.3 Joint spike–LFP inference

We fuse LFP and spike observations in a single linear–Gaussian state-space model. Fix a frequency *ω*_*j*_ and work on the spike-bin grid 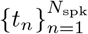 with spacing Δ_spk_. To apply standard Kalman filtering, we represent the *M* complex taper-specific latent states in real form by stacking real and imaginary parts:

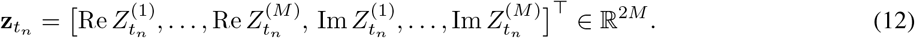

Under our convention *N*_ℂ_ (*µ, q*) with *q* = E |*X* − *µ*| ^2^, the equivalent real-valued representation has Var(Re *X*) = Var(Im *X*) = *q/*2.

The OU dynamics discretized at Δ_spk_ give

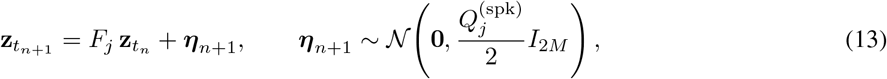

where 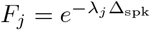 and 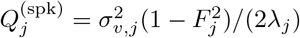.

Two types of observations attach to this latent process. At block centers *t*_*k*_, the LFP provides a 2*M*-dimensional Gaussian observation. Stacking real and imaginary parts of the multitaper coefficients into 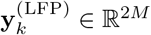, we have

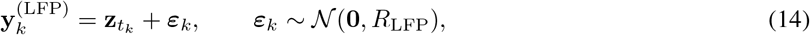

where 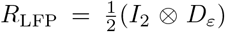 with 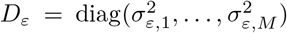. At every spike bin, the pseudo-observation (11) contributes a scalar Gaussian row. The linear predictor *ψ*_*n*_ depends on 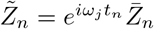, which is a known linear function of 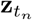. Subtracting the intercept and spike-history terms from (11) yields

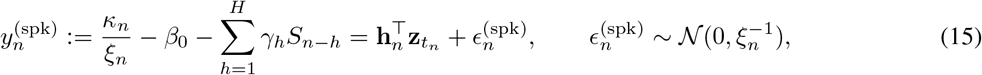

where **h**_*n*_ ∈ ℝ^2*M*^ encodes the taper averaging and phase rotation. With 1_*M*_ denoting the all-ones vector in ℝ^*M*^ and *θ*_*n*_ = *ω*_*j*_*t*_*n*_,

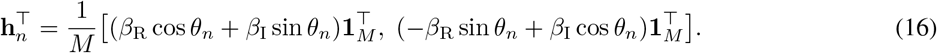

The derivation of this observation vector from the phase-rotated taper average appears in Appendix B.2.

The inference problem is now a standard linear–Gaussian state-space model on the spike-bin grid. At each time *t*_*n*_, we apply the prediction step (13). At spike bins, we apply the scalar observation update (15). At the subset of bins corresponding to LFP block centers, we additionally apply the vector observation update (14). A forward Kalman filter and backward Rauch–Tung–Striebel smoother yield posterior means 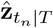 and covariances 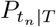 at every spike bin. A simulation smoother [Carter and Kohn, 1994, Frühwirth-Schnatter, 1994] produces joint draws of the entire latent trajectory. Details of the forward filtering, backward sampling algorithm appear in Appendix B.1.

We estimate the model parameters by iterating between three blocks. First, given the current latent trajectory (or its smoothed moments), we draw the Pólya–Gamma auxiliary variables *ξ*_*n*_ ~ PG(1, |*ψ*_*n*_|) for each spike bin. Second, given *{ξ*_*n*_*}* and the latent trajectory, the coupling coefficients ***β*** = (*β*_0_, *β*_R_, *β*_I_, *γ*_1_, …, *γ*_*H*_)^⊤^ have a Gaussian conditional posterior. Defining the design matrix **D** with rows 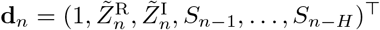 and the diagonal precision matrix 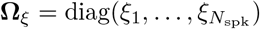, the posterior is

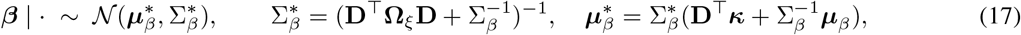

where 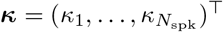 and (***µ***_*β*_, ∑_*β*_) is a Gaussian prior. When using smoothed moments rather than sampled trajectories, the required expectations are computed via the linear transformation in Appendix B.2. Third, given *ξ*_*n*_ and ***β***, we update the latent trajectory via the Kalman smoother described above, and update the OU parameters 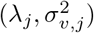 and observation noise 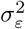 by EM using smoothed sufficient statistics (Appendix B.1). The full procedure is summarized in Algorithm 1.

#### Algorithm 1

Joint inference at frequency *ω*_*j*_ (one outer iteration)

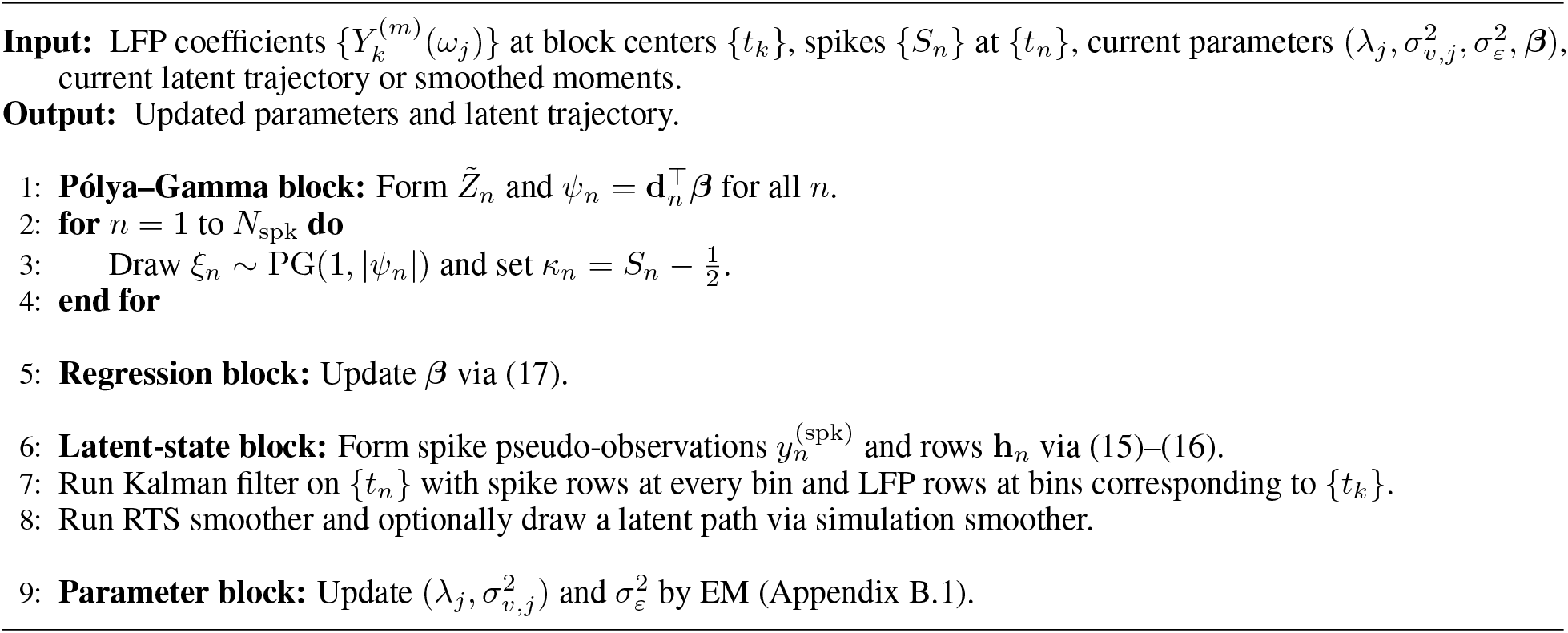

### 2.4 Hierarchical model for multi-trial data

When multiple trials are available, we decompose the latent spectral trajectory into a shared component *X* that is common across trials and trial-specific deviations *δ*_*r*_:

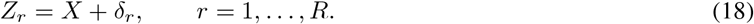

Both *X* and *δ*_*r*_ follow independent Ornstein–Uhlenbeck dynamics with potentially different timescales. The shared process evolves with parameters 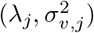, while the deviations evolve with 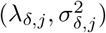. This structure captures the intuition that neural oscillations have a stereotyped component that is consistent across trials, plus trial-to-trial variability that may reflect fluctuations in attention, arousal, or other latent factors.

Inference proceeds via a two-pass approximation. The first pass pools observations across trials using precision weighting to estimate the shared trajectory 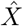. Pooling leverages the fact that E[*δ*_*r*_] = 0, so the cross-trial average isolates the shared component. The second pass estimates each trial’s full trajectory 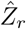 using per-trial observations, from which we recover the deviation as 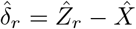. Both passes use the joint Kalman filtering framework described in Section 2.3, with Pólya–Gamma augmentation to incorporate spike observations. Details of the pooling formulas and parameter updates appear in Appendix C.

## 3 Applications: single trial

### 3.1 Ground-truth simulation

We evaluated the inference framework on simulated data where all generative parameters are known. The simulation comprises a 300-second recording with LFP sampled at 1000 Hz and spike trains binned at 1 ms resolution. Figure 1(a) displays a snippet for the LFP signal and spike trains. The LFP contains oscillatory components at six frequencies. Four coupled bands (11, 19, 27, and 43 Hz) that drive spike timing in a subset of units, and two uncoupled bands (7 and 35 Hz) that carry oscillatory signal but do not modulate spiking. All other frequencies in the 1–60 Hz range contain only observation noise. Five simulated units each couple to a random subset of three coupled bands, with coupling strengths |*β*_*C*_| drawn uniformly from a specified range and preferred phases drawn uniformly on (−*π, π*]. Simulation details appear in Section D.

Figure 1(c) compares spectrogram estimates across methods. Standard multitaper analysis captures dynamics in the signal band but is overwhelmed by observation noise. The continuous-time state-space multitaper model (CT-SSMT), using LFP data alone, exploits temporal autocorrelation to denoise the spectrogram. Frequency bands with true oscillatory signal exhibit smooth amplitude trajectories, while bands containing only observation noise are regularized toward zero. This denoising arises because the EM algorithm infers large *λ*_*j*_ for noise bands, causing the Kalman smoother to shrink estimates toward the prior mean.

Incorporating spike train data further refines amplitude estimates at coupled frequencies. Panel (d) displays time-resolved amplitude trajectories at 11 Hz and 27 Hz for two representative time windows. The Joint SSMT estimate (orange) tracks the ground-truth amplitude (dashed black) more closely than either multitaper (blue) or LFP-only CT-SSMT (magenta), with 95% credible intervals quantifying estimation uncertainty. Panel (e) summarizes this improvement by computing Pearson correlations between estimated and true power in non-overlapping 20-second windows. At coupled frequencies (11, 19, 27, 43 Hz), Joint SSMT model achieves systematically higher correlations than either comparison method. At uncoupled signal frequencies (7, 35 Hz), spike trains do not enter joint inference because no coupling was detected throughout the inference procedure.

Joint SSMT simultaneously estimates spike–field coupling coefficients with uncertainty quantification. Figure 2(a) displays posterior samples of the complex coupling coefficient *β*_*C*_ = *β*_*R*_ + *iβ*_*I*_ for Unit 0 at six frequencies. At truly coupled frequencies (11, 19, 27 Hz for this unit), the posterior mean lies away from the origin and the 95% credible ellipse excludes zero. At uncoupled frequencies (7, 35, 43 Hz), the posterior concentrates near the origin.

**Figure 2:**
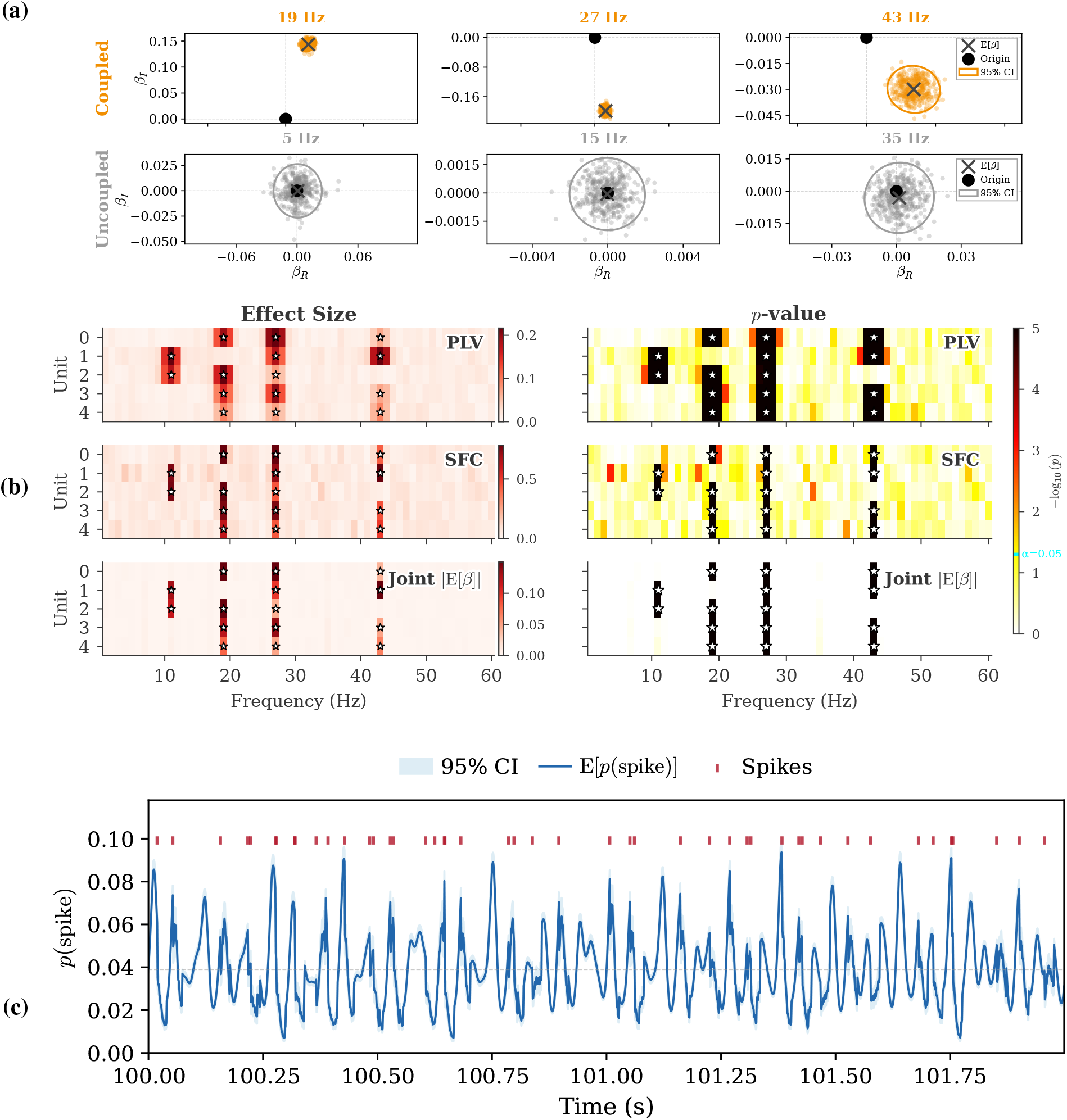
Coupling detection via Joint SSMT. **(a)** Posterior samples of complex coupling coefficients *β*_*C*_ = *β*_*R*_ + *iβ*_*I*_. Top row: coupled frequencies show posteriors displaced from the origin with 95% credible ellipses excluding zero. Bottom row: uncoupled frequencies show posteriors centered near the origin. **(b)** Effect size (left) and − log_10_(*p*) (right) heatmaps comparing PLV, SFC, and Joint SSMT. White stars indicate true couplings. Joint SSMT shows sharper frequency selectivity with fewer false positives. **(c)** Posterior spike probability for a representative unit under Joint SSMT. The mean (blue) and 95% credible interval (shaded) show oscillatory modulation of spiking propensity. Red ticks mark observed spikes, which occur preferentially during high-probability phases.

We assess statistical significance using the Wald test. Let 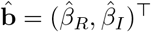 denote the posterior mean and let *V*_*β*_ denote the 2 × 2 posterior covariance matrix. The Wald statistic is

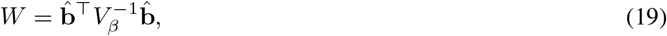

which follows a 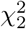 distribution under the null hypothesis *β*_*C*_ = 0. We also considered a phase concentration test analogous to the Rayleigh test used for PLV, but found the Wald test more reliable in finite samples.

Figure 2(b) compares effect sizes and *p*-values across methods. PLV and SFC detect coupling at the correct frequencies but exhibit spectral leakage, reporting elevated values at neighboring frequencies. The parametric significance tests available for PLV (Rayleigh test) and SFC (coherence *F*-test) fail to control false positives at the nominal level. Many uncoupled unit–frequency pairs exceed the *α* = 0.05 threshold. The Wald test applied to the joint model posterior identifies only the truly coupled pairs, with no false positives among the 25 uncoupled combinations.

Beyond coupling detection, the joint model yields a complete posterior over the instantaneous spike probability *σ*(*ψ*_*n*_) at each time bin. Figure 2(c) displays this posterior for a representative unit. The mean spike probability (blue) and 95% credible interval track the oscillatory modulation of firing propensity, and observed spikes (red ticks) occur preferentially during high-probability phases.

### 3.2 Propofol-induced anesthesia

We applied the method to multielectrode recordings from macaque cortex during propofol-induced loss of consciousness, a dataset previously analyzed in Bastos et al. [2021]. LFP and single-unit activity were recorded from ventrolateral prefrontal cortex (vlPFC) and posterior parietal area 7b while propofol was infused intravenously. The recording was partitioned into six epochs, namely Baseline (pre-drug), Drug Onset, two periods of loss of consciousness (LOC 1, LOC 2), Drug Stop, and Recovery. Surgical, pharmacological, and experimental details appear in Bastos et al. [2021].

Figure 3(a) displays raw LFP and spike rasters during Baseline, LOC 1, and Recovery. During loss of consciousness, the LFP exhibits prominent slow oscillations and spiking becomes sparse, with spikes clustering at particular phases of the slow rhythm. The spike-triggered average in panel (b) confirms this observation. During LOC epochs, the peri-spike LFP shows a pronounced oscillatory deflection absent during Baseline and Recovery.

**Figure 3:**
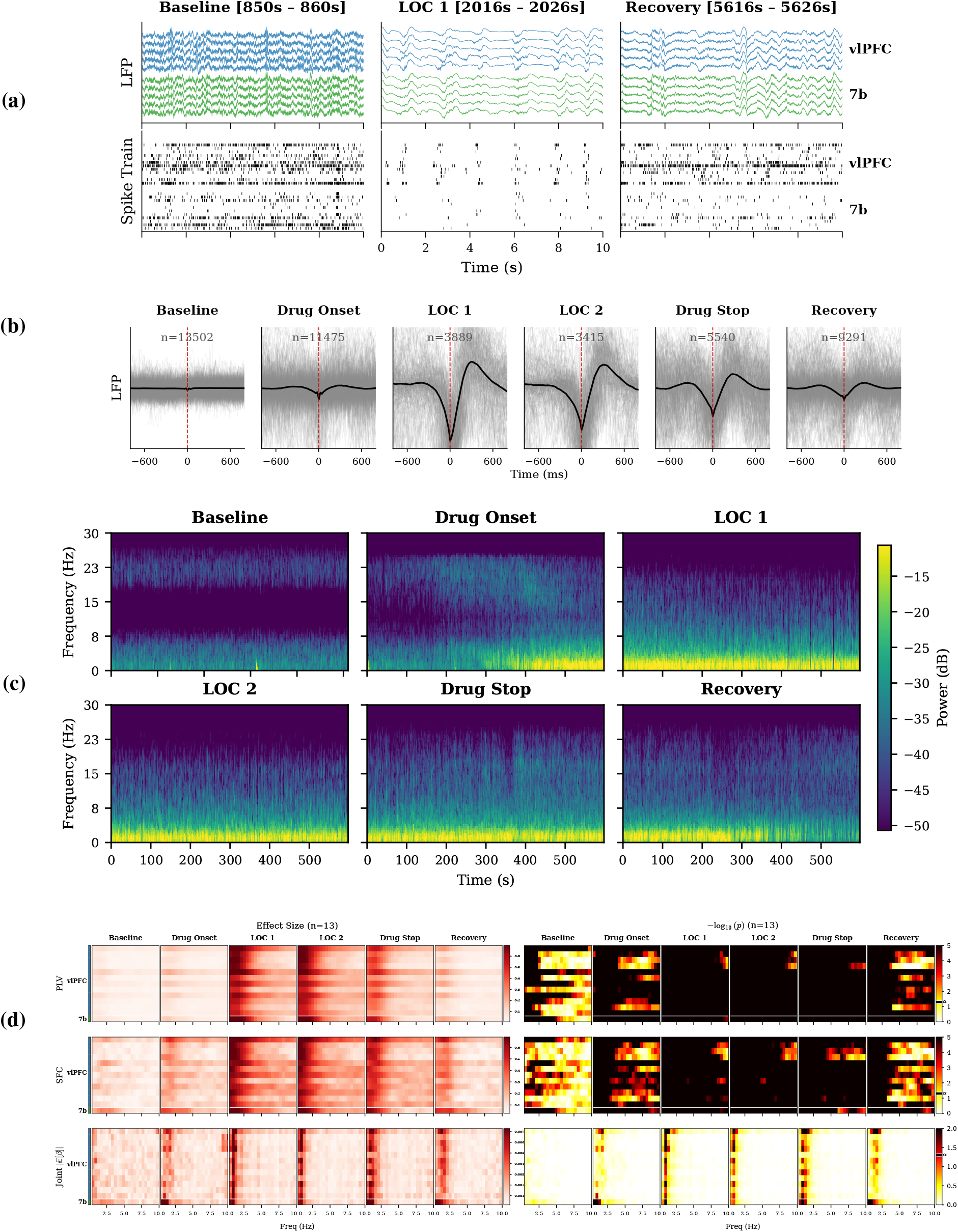
Joint SSMT on anesthesia data. **(a)** Raw LFP and spikes during baseline, LOC, and recovery. **(b)** STA reveals enhanced phase-locking during anesthesia. **(c)** Spectrogram showing increased low-frequency power during LOC. **(d)** Coupling effect sizes (left) and significance (right) for PLV, SFC, and the Joint SSMT Wald test. Each row corresponds to a unit recorded from the same electrode as the LFP. Joint SSMT identifies coupling at a specific slow-oscillation frequency near 0.8 Hz.

Panel (c) presents Joint SSMT spectrograms for each epoch. The inferred spectrogram reveals increased low-frequency power during LOC, consistent with the slow oscillations characteristic of propofol anesthesia. Compared to the standard multitaper spectrogram (Figure S1), the state-space estimate provides clearer delineation of spectral structure.

Panel (d) compares coupling effect sizes and significance across methods. PLV and SFC both indicate elevated coupling in the low-frequency range during LOC epochs, but this coupling is spectrally diffuse, spanning roughly 0.4–5 Hz with no clear peak. The joint model identifies coupling at a specific slow-oscillation frequency near 0.8 Hz, consistent with the intuitive observations from spike trigger averages.

Parametric tests for PLV (Rayleigh) and SFC (coherence *F*-test) declare nearly all low-frequency bins significant during LOC, providing little discrimination. We also computed non-parametric *p*-values via circular shifting and spike-train jittering (Figure S2). These permutation tests similarly yield a diffuse pattern across the low-frequency range. The Wald test applied to the joint model posterior isolates a narrow frequency band with significant coupling, consistent with entrainment of spiking to a specific slow-oscillation rhythm rather than broadband low-frequency modulation.

## 4 Application to data with trial structure

### 4.1 Ground-truth simulation

To test the hierarchical framework, we simulated 100 trials of 10 seconds each. Six frequencies carry true oscillatory signal. Four (11, 19, 27, 43 Hz) drive spike timing, while two (7, 35 Hz) contain oscillatory power but no spike coupling. The hierarchical decomposition *Z*_*r*_ = *X* + *δ*_*r*_ partitions each trial’s spectral trajectory into a shared component *X* and a trial-specific deviation *δ*_*r*_. We set the shared process to evolve slowly (half-bandwidth 0.05 Hz) and the deviations to decay fourfold faster. Five units couple to distinct subsets of the four coupled bands. Coupling magnitudes and preferred phases vary across units. Full simulation parameters appear in Appendix E.1.

Figure 4(a) shows trial-averaged spectrograms. The hierarchical model attenuates noise-only frequencies while preserving all six signal bands, whether or not they couple to spikes. The central question for trial-structured data is whether the method captures trial-specific fluctuations encoded in *δ*_*r*_, not merely the shared template *X*. Panel (c) displays amplitude trajectories at 11 and 19 Hz across four individual trials. Joint SSMT mean estimate with credible intervals follows fluctuations in ground-truth amplitude that differ from trial to trial. Panel (b) quantifies correlations between estimated and ground-truth power for each trial. At coupled frequencies, Joint SSMT is substantially higher than multitaper or LFP-only inference.

**Figure 4:**
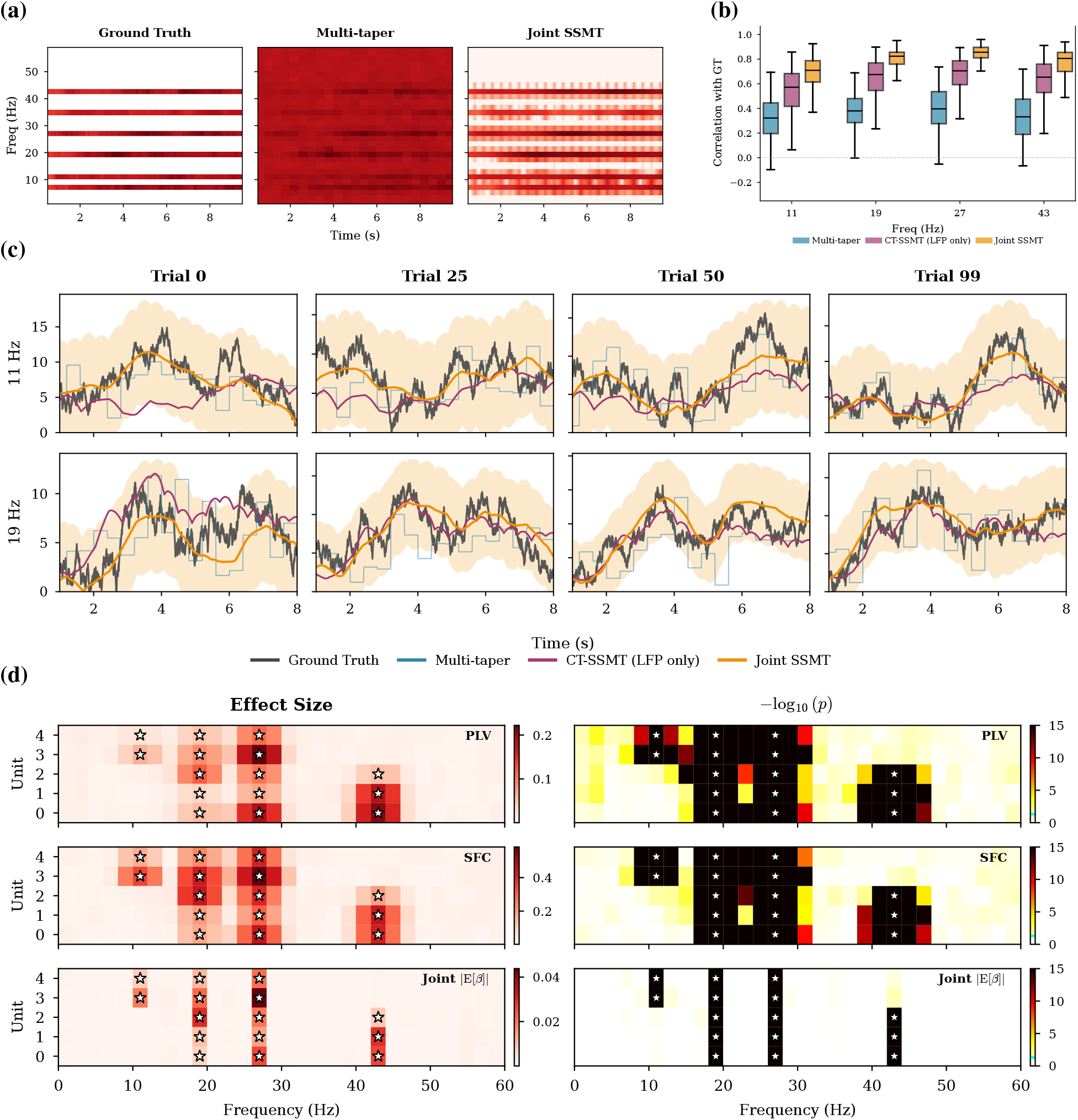
Trial-structured spike–field coupling inference. **(a)** Trial-averaged spectrograms: ground truth, multitaper, and Joint SSMT. **(b)** Correlation with ground truth. Joint SSMT (orange) achieves near-perfect correlation. **(c)** Single-trial dynamics at 11 and 19 Hz with 95% CI. **(d)** Effect sizes (left) and statistical significance as −log_10_(*p*) (right) for PLV, SFC, and Joint SSMT. Each row corresponds to a simulated unit, and stars indicate ground-truth coupled frequency bands.

Panel (d) compares coupling detection. Stars mark ground-truth couplings. PLV and SFC show elevated effect sizes at bands with coupling, but their associated parametric tests declare many other coupling pairs significant. By contrast, the Wald test on the joint posterior detects coupling exclusively at the four truly coupled frequencies, with no false positives at signal-only bands or among uncoupled unit–frequency pairs.

### 4.2 Associative learning task

We applied the method to multielectrode recordings from macaque hippocampus and prefrontal cortex during a sample-sample paired associates (SSPA) task. On each trial, the animal viewed a sample stimulus, retained it through a delay period, and then indicated whether a test stimulus matched or did not match a learned associate. The task requires both recognition of individual stimuli and retrieval of learned associations, engaging both hippocampal and prefrontal circuits.

LFP and single-unit activity were recorded from multiple brain areas including CA3, ventral prefrontal cortex (PFCv), subiculum (Sub), and the head of the caudate nucleus (hCd). We analyzed the last 400 valid trials, pooled across match/nonmatch and correct/incorrect conditions to maximize statistical power. The analysis window spanned −0.5 to 3.5 seconds relative to sample onset, covering baseline, stimulus presentation, delay, and response epochs. The frequency grid ranged from 1 to 80 Hz in 2 Hz steps, with 300 ms multitaper windows. We included all units with mean firing rates exceeding 1 Hz. Full analysis details appear in Appendix E.2.

Figure 5(a) displays the Joint SSMT inference results for 36 units sorted by brain area. Several units in PFCv and hCd exhibit clear coupling to delta/theta (1–8 Hz) and beta (12–30 Hz) frequency bands, with *p* < 0.001. The coupling is frequency-specific. Elevated effect sizes concentrate in narrow bands rather than spanning broad frequency ranges.

**Figure 5:**
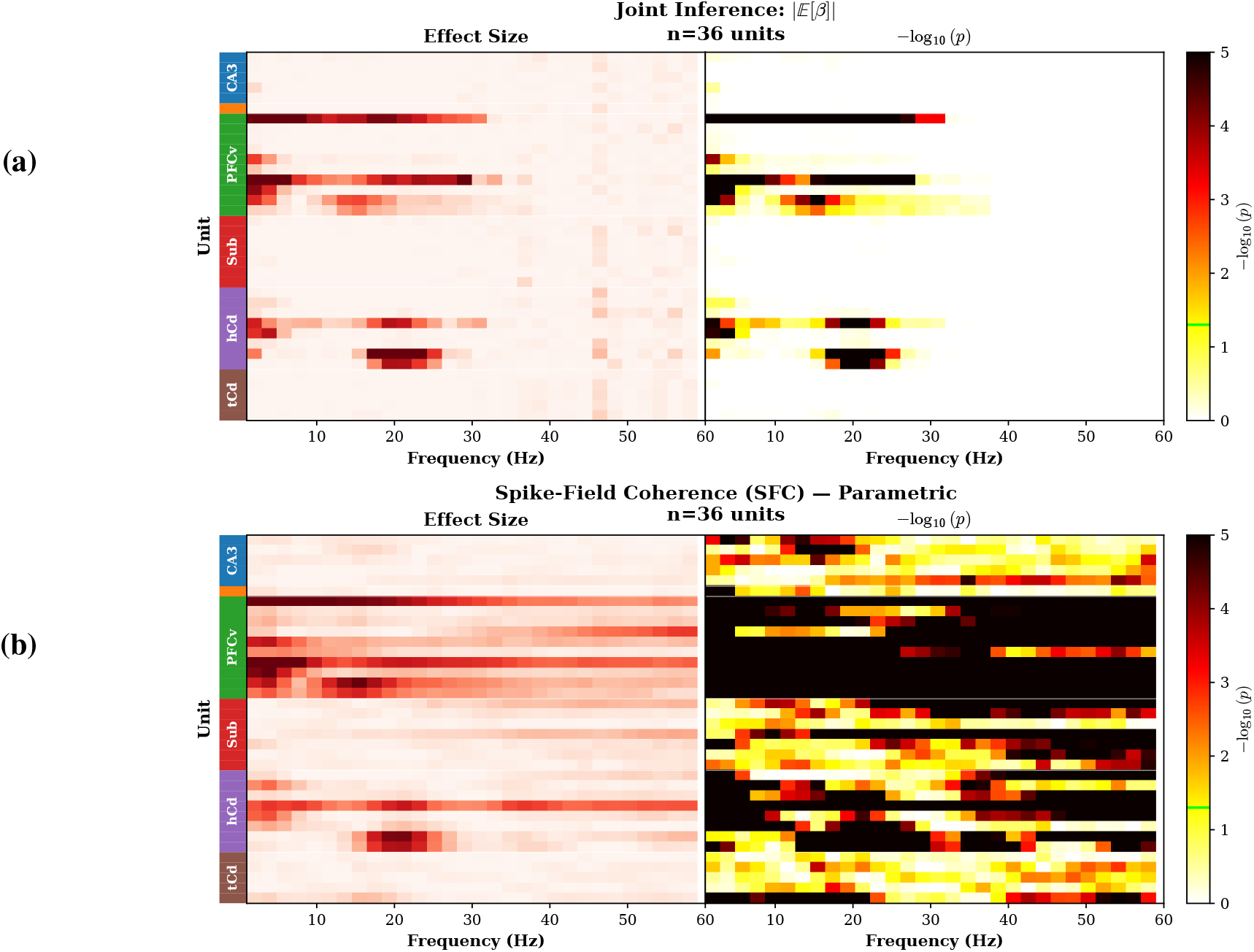
Association learning task: trial structure and spectro-temporal signatures. **(a)** Population-level spike– field coupling from Joint SSMT: effect size |E[*β*_*C*_]| and −log_10_(*p*), sorted by area. **(b)** Coupling effect sizes and p-values for SFC (parametric test). Joint SSMT identifies more specific spike–field coupling frequency bands and gives meaningful uncertainty quantification.

Panel (b) shows the corresponding SFC analysis with parametric significance testing. While SFC detects some of the same coupled pairs, the effect sizes are more diffuse across frequencies, reflecting the spectral leakage inherent in coherence estimation. The parametric F-test declares many unit–frequency pairs significant, including some at frequencies where the joint model finds no evidence of coupling.

Supplementary Figures S4 and S5 present additional comparisons using PLV with the Rayleigh test and permutation-based significance testing for both PLV and SFC. The permutation tests for PLV and SFC also do not meaningfully yield frequency-specific significance detection. Across all comparisons, the Wald test applied to the joint posterior provides the sharpest frequency discrimination, identifying coupling in specific bands implicated in associative learning Brincat and Miller [2015].

## 5 Connection to classical spike-field coupling measures

The generative model in Section 2.1 provides a statistical framework within which classical spike-field coupling measures can be expressed in closed form. We derive these expressions for the phase-locking value (PLV) and spike-field coherence (SFC). The resulting formulas make explicit how PLV and SFC depend on model parameters, including the baseline log-odds *β*_0_, coupling strength |*β*_*C*_|, and OU bandwidth *λ*, offering insights into the behavior of these measures that are not immediately apparent from their empirical definitions. For the analytic derivations, we set the spike-history terms to zero (*γ*_*h*_ = 0) and assume the OU process is in stationarity.

### 5.1 Phase-locking value

The phase-locking value (PLV) measures the concentration of spike times around a preferred signal phase. We derive its theoretical form under the logistic spike model.

#### Proposition 1

(PLV for the logistic spike model). *Let Z* ~ *CN* (0, *q*) *and S* | *Z* ~ Bernoulli(*σ*(*β*_0_ + *β*_R_ Re(*Z*) + *β*_I_ Im(*Z*))). *Define* 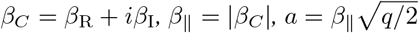, *and let* 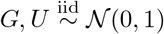. *Then*

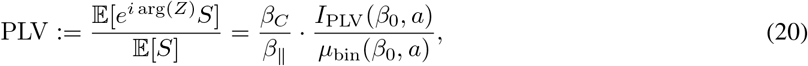

*where µ*_bin_(*β*_0_, *a*) = E_*G*_[*σ*(*β*_0_ + *aG*)] *and I*_PLV_(*β*_0_, *a*) = E_*G*_[*σ*(*β*_0_ + *aG*) · *G* · *h*(*G*)] *with*

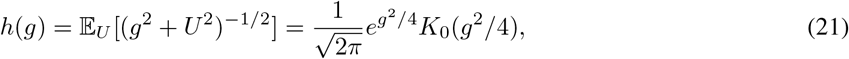

*and K*_0_(·) *is the modified Bessel function of the second kind. In particular*, arg(PLV) = arg(*β*_*C*_).

The proof rotates coordinates to align with the coupling direction *β*_*C*_, reducing the problem to a one-dimensional integral that can be evaluated via iterated expectations. Details appear in Appendix F.1.

#### Corollary 1

(PLV for the Joint SSMT spike-field coupling model). *Consider the spike-field coupling model* (6). *Let* 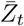 *follow the OU process* (3) *with stationary variance* 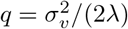, *and let* 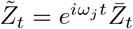 *be the phase-rotated spectral coefficient at frequency ω*_*j*_. *The empirical PLV, computed as*

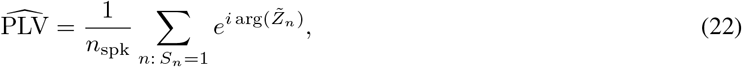

*converges almost surely to the theoretical value* (20) *as observation time grows. The PLV magnitude depends on* (*λ, σ*_*v*_) *only through q, and is independent of the frequency ω*_*j*_.

The corollary follows from the circular symmetry of the complex Gaussian distribution, which ensures that phase rotation preserves the marginal 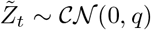, combined with the ergodicity of the OU process. The frequency *ω*_*j*_ enters only through the deterministic rotation and does not affect the marginal variance *q* or any quantity in the PLV formula.

Figure 6 validates these results. Panel (a) shows PLV as a function of coupling strength |*β*_*C*_| for several values of the baseline log-odds *β*_0_. Stronger coupling yields higher phase-locking, with lower firing rates (more negative *β*_0_) producing larger PLV at fixed coupling strength. Panel (b) displays PLV versus *β*_0_ for several coupling strengths. PLV decreases monotonically with increasing *β*_0_, reflecting the reduced sensitivity of the sigmoid nonlinearity at high firing rates. Panels (a) and (b) validate the proposition by sampling *Z* i.i.d. from *CN*(0, *q*). Panel (c) validates the Corollary by simulating the full OU dynamics with phase rotation at multiple frequencies and decay rates. All curves collapse onto a single theoretical prediction, confirming that PLV depends on (*λ, σ*_*v*_) only through the stationary variance *q* and is independent of the carrier frequency *ω*_*j*_.

**Figure 6:**
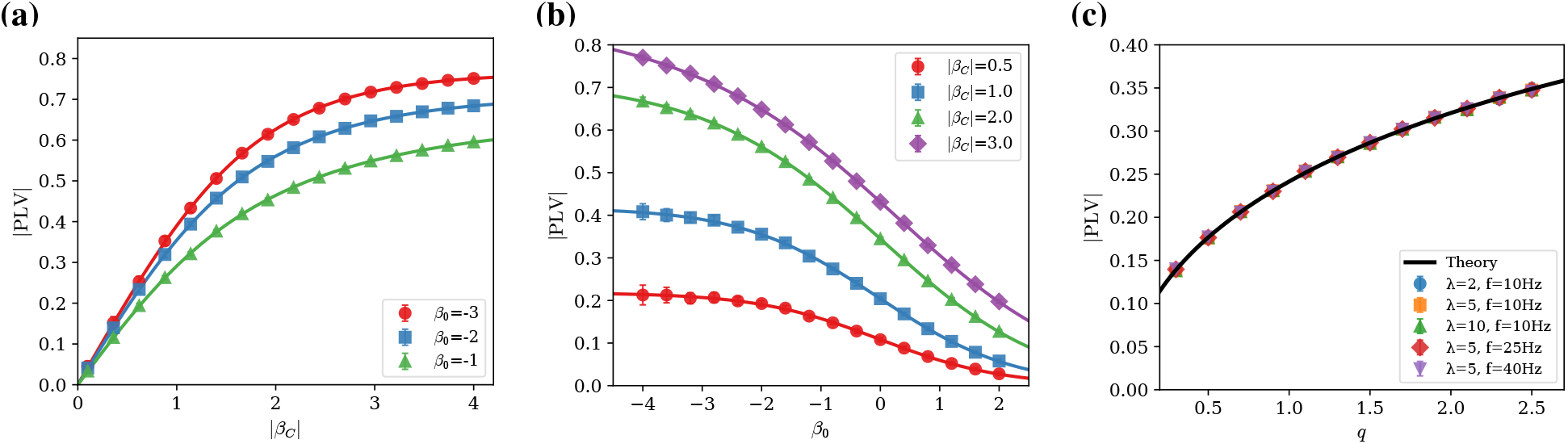
Validation of closed-form phase-locking value. **(a)** PLV versus coupling strength |*β*_*C*_| for different *β*_0_; lower firing rates yield higher PLV at fixed coupling. **(b)** PLV versus *β*_0_ for different |*β*_*C*_|; PLV decreases with increasing firing rate. **(c)** PLV versus *q* for different (*λ, f*_0_) combinations; all curves collapse, confirming independence from *λ* and *ω*_*j*_. Solid curves: theory; markers: Monte Carlo (*±*1 s.d.).

The closed-form PLV reveals that phase-locking depends not only on coupling strength |*β*_*C*_| but also on the baseline firing rate through *β*_0_. At fixed coupling, a unit with lower firing rate (more negative *β*_0_) exhibits higher PLV, because the sigmoid nonlinearity is more sensitive in its low-rate regime. This interaction means that changes in firing rate across experimental conditions, for instance during anesthesia (Section 3.2), can alter the empirical PLV even when the underlying coupling has not changed. The joint model estimates *β*_0_ and *β*_*C*_ separately, allowing these effects to be disentangled.

### 5.2 Spike-field coherence

We now derive a closed-form expression for the spike-field coherence (SFC) under the joint model. For the analytic LFP signal 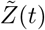 and spike train *S*(*t*), the coherence at frequency *f* is defined as

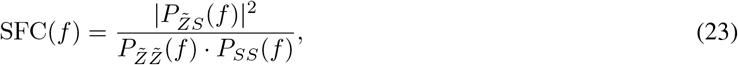

where 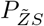 denotes the cross-spectral density and 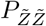, *P*_*SS*_ denote the auto-spectral densities. This quantity measures the fraction of spike train power at frequency *f* that is linearly predictable from the LFP.

The derivation requires two intermediate results. The first establishes the cross-spectrum between the LFP and the spike train.

#### Proposition 2

(Cross-Spectrum). *Under the model* 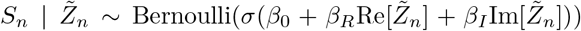, *the cross-spectral density satisfies*

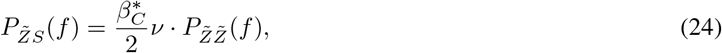

*where β*_*C*_ = *β*_*R*_ + *iβ*_*I*_ *and* ν = E[*σ*(*β*_0_ + *aG*)(1 − *σ*(*β*_0_ + *aG*))] *with* 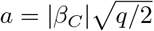 *and G* ~ *N* (0, 1).

The cross-spectrum is proportional to the LFP spectrum, with a constant of proportionality determined by the coupling coefficient and the expected conditional variance ν of the Bernoulli process. The result follows from Stein’s lemma applied to the logistic nonlinearity. The proof appears in Appendix F.2.

The second result decomposes the spike train spectrum into signal and noise components.

#### Proposition 3

(Approximate spike spectrum). *The spike train power spectral density at the carrier frequency is*

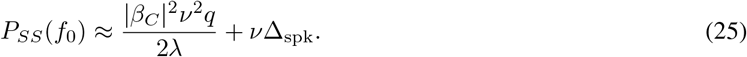

The formula admits interpretation as a signal-to-noise ratio. The term |*β*_*C*_| ^2^ν*q/*(2*λ*) in the denominator represents the signal power from coherent rate modulation, while νΔ_spk_ represents the incoherent Bernoulli noise. When signal dominates, SFC → 1. When noise dominates, SFC → 0.

Combining these results yields the main result.

#### Proposition 4

(Closed-Form SFC). *At the carrier frequency f*_0_, *the spike-field coherence is*

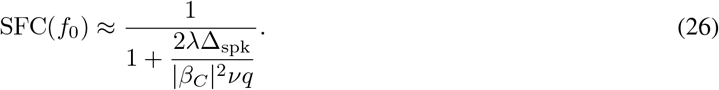

The closed-form expression reveals a structural difference between SFC and PLV. Whereas PLV depends on the model parameters only through the combination 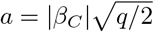, SFC depends separately on *q* and the OU decay rate *λ*. This difference arises because SFC involves the LFP power spectrum 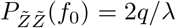, which depends on the ratio *q/λ*, not merely on *q*. Increasing *λ* at fixed *q* broadens the spectral bandwidth, reducing peak power at *f*_0_ while leaving the Bernoulli noise floor unchanged, thereby decreasing SFC. Figure 7 validates the closed-form expression against Monte Carlo simulation. Panel (a) shows SFC as a function of coupling strength |*β*_*C*_| for several values of the baseline log-odds *β*_0_. Stronger coupling yields higher coherence, with the effect modulated by firing rate. Panel (b) displays SFC versus *β*_0_ for several coupling strengths. SFC has non-monotonic dependency on *β*_0_. Panel (c) plots SFC against the stationary variance *q* for different combinations of (*λ, f*_0_). In contrast to the PLV validation (Figure 6c), where curves for different *λ* collapse when plotted against *q*, the SFC curves remain separated, reflecting the explicit *λ*-dependence in the formula.

**Figure 7:**
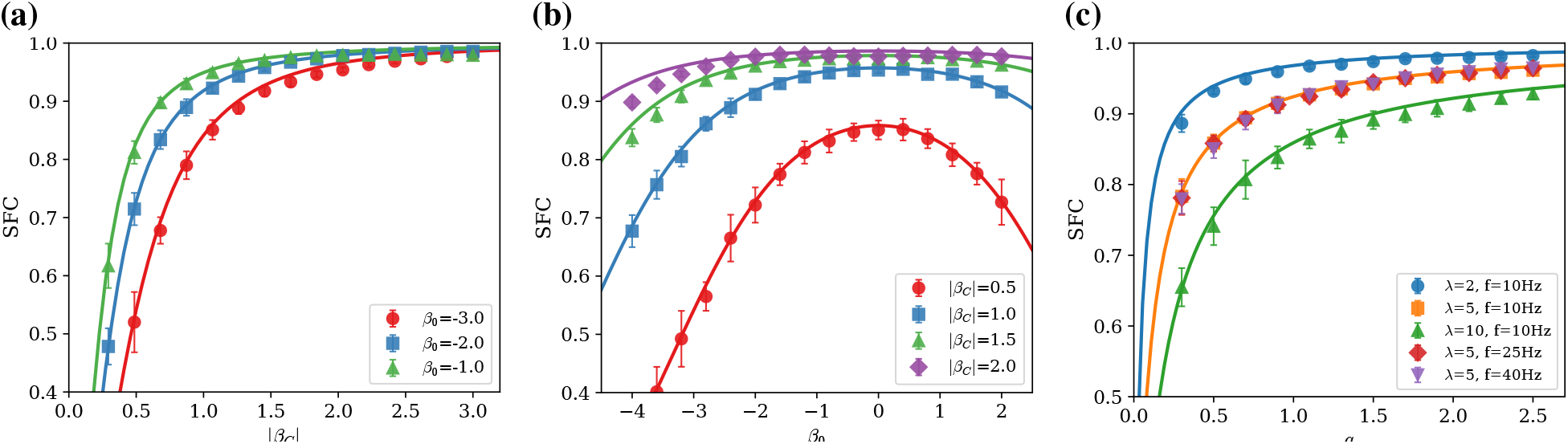
Validation of closed-form spike–field coherence. **(a)** SFC versus coupling strength |*β*_*C*_| for different *β*_0_. (**b**) SFC versus *β*_0_ for different |*β*_*C*_|. **(c)** SFC versus *q* for different (*λ, f*_0_) combinations; curves with the same *λ* overlap regardless of carrier frequency. Solid curves: theory; markers: Monte Carlo (*±*1 s.d.).

Together, the closed-form expressions for PLV and SFC characterize how each measure responds to the parameters of the generative model. Both depend on *β*_0_, but in qualitatively different ways. PLV decreases monotonically with increasing firing rate, while SFC exhibits a non-monotonic dependence (Figure 7b), first increasing and then decreasing as *β*_0_ grows. These observations suggest that empirical changes in PLV or SFC across conditions may partly reflect changes in firing rate rather than in the coupling itself. The generative model approach estimates all parameters jointly, providing a more complete account of the spike-field relationship than either summary statistic alone.

## 6 Discussion

We have developed **Joint SSMT**, a Bayesian state-space framework that jointly infers time-varying spectrograms and point-process coupling. Joint SSMT extends the state-space multitaper paradigm of Kim et al. Kim et al. [2018] by incorporating discrete event observations through Pólya–Gamma augmentation Polson et al. [2013]. Multitaper Fourier coefficients and spike observations share a common latent spectral process, enabling joint inference of oscillatory dynamics and spike-field coupling. LFP measurements constrain spectral dynamics at coarse temporal resolution while spike times refine estimates at millisecond precision. Simulation studies with known ground truth demonstrate that the framework recovers coupling coefficients with principled uncertainty quantification and localizes coupling to narrower frequency bands than classical PLV or SFC. We applied the method to two experimental datasets. In propofol-induced anesthesia recordings, the framework identifies spike-field coupling consistent with previous reports Bastos et al. [2021] but with finer frequency resolution. In an associative learning task, the method reveals unit-specific theta- and beta-band coupling in prefrontal cortex and hippocampus. The closed-form relationships derived in propositions 1 and 4 express PLV and SFC as functions of the generative model parameters, revealing their distinct dependencies on firing rate, coupling strength, and spectral dynamics.

### 6.1 Computational implementation

The inference algorithm exploits conditional independence structure for efficient computation. Each frequency band admits an independent Kalman filter, and the Pólya-Gamma augmentation renders the spike likelihood conditionally Gaussian, avoiding costly Metropolis-Hastings updates. We implemented the sampler in JAX, which enables just-in-time compilation and automatic vectorization across frequency bands, tapers, and units. The Pólya-Gamma random variates are drawn using the method of Windle et al. Windle et al. [2014], implemented in JAX to enable GPU parallelization. On a single GPU, inference for the trial-structured simulation (100 trials, 10 seconds each, 30 frequency bands, 5 units) completes in under 5 minutes. These runtimes permit application to large datasets with more recorded units.

### 6.2 Related analyses

Beyond the analyses demonstrated in this paper, the Joint SSMT framework supports several direct analyses with minimal additional modeling. Joint SSMT inherits several useful properties from the state-space multitaper paradigm of Kim et al. Kim et al. [2018]. The Kalman smoother provides not only spectrogram estimates but also time-domain signal extraction. Given the smoothed increment differences 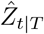 at each frequency, the denoised LFP can be reconstructed by inverse Fourier transform, and selective filtering (low-pass, high-pass, band-pass) can be performed by zeroing increment differences outside the desired frequency range. Instantaneous amplitude and phase at any frequency follow directly from the complex-valued latent state, with posterior means and variances that incorporate spike information.

Cross-spectral analysis can be incorporated into the joint framework. Given two LFP signals *Y* ^(1)^ and *Y* ^(2)^ recorded from different electrodes or brain regions, one can define a bivariate latent spectral process and estimate the cross-spectrogram *P*_12_(*f, t*) via the smoothed cross-products of the increment differences. Coherence between regions can then be computed as a function of time, with uncertainty quantified through the posterior distribution. Incorporating spike observations from neurons in each region would yield a joint model of inter-regional LFP coherence and spike-field coupling, enabling investigation of how oscillatory coordination between areas relates to single-neuron phase-locking.

Multiple LFP channels can be incorporated into a single inference by augmenting the observation model. If *C* channels share a common latent spectral process (for example, electrodes within a local region), the multitaper observations from all channels can be stacked into a single observation vector at each block center. The observation noise covariance becomes block-diagonal across channels, and the Kalman filter proceeds as before with a larger observation dimension. Alternatively, if the scientific question concerns differences between channels, a hierarchical model analogous to the trial-structured extension can decompose the latent state into shared and channel-specific components.

The Pólya–Gamma augmentation strategy applies not only to Bernoulli likelihoods but to binomial likelihoods of the form *S* ~ Binomial(*L, p*) with logit(*p*) = *ψ* Polson et al. [2013]. This observation permits coarse temporal binning of spike counts. If spike times are aggregated into bins of width Δ_*bin*_ > Δ_spk_, the count *S*_*b*_ ∈ {0, 1, …, *L*_*b*_} in bin *b* follows a binomial distribution with *L*_*b*_ trials (the number of fine bins within the coarse bin) and success probability *σ*(*ψ*_*b*_). The augmented likelihood involves drawing *ξ*_*b*_ ~ PG(*L*_*b*_, |*ψ*_*b*_|), and inference proceeds as before. Coarse binning reduces the state dimension and may improve computational efficiency when millisecond-resolution coupling is not required.

### 6.3 Theoretical and methodological generalizations

We next describe extensions that modify the latent dynamics or coupling structure and therefore require additional modeling and inference. Several assumptions of the current model could be relaxed to accommodate more complex data structures. The Ornstein-Uhlenbeck dynamics impose exponential autocorrelation on the latent spectral process, which is appropriate for narrowband oscillations with stable center frequency. Transient spectral events—bursts, chirps, or abrupt frequency shifts—violate this assumption. One remedy is to replace the OU process with a switching state-space model in which the latent state can transition between multiple dynamical regimes Linderman et al. [2016]. Each regime would have its own OU parameters, and the posterior distribution over regime sequences would identify when spectral transitions occur.

The logistic link function models sinusoidal modulation of firing probability. The linear predictor 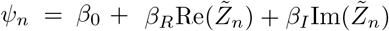 implies that the log-odds of spiking varies as |*β*_*C*_| cos(*ω*_0_*t* − *ϕ*) where *ϕ* = arg(*β*_*C*_). This formulation captures single-peaked phase tuning within each band, but it does not capture multi-peaked phase preferences or waveform-shape sensitivity (e.g., distinct responses to rising versus falling flanks) without adding higher harmonics or richer phase/amplitude basis functions. Generalized additive models that replace the linear predictor with smooth functions of phase and amplitude could capture such effects. Basis function expansions—circular splines for phase, monotone splines for amplitude—would maintain interpretability while providing additional flexibility. The cost is increased model complexity and potential overfitting with limited data.

The current spike model includes a history kernel to capture refractory effects and short-term adaptation, but it treats each neuron independently. Population models that include coupling between neurons could reveal how oscillatory coordination emerges from network interactions. A natural extension is to add terms of the form ∑_ℓ≠*i*_ *w*_*i*ℓ_*S*_ℓ_(*t* −*τ*) to the linear predictor for neuron *i*, representing excitatory or inhibitory interactions from other neurons in the ensemble. The coupling weights *w*_*il*_ could be estimated jointly with the spike-field coupling coefficients, yielding a unified model of field-to-spike and spike-to-spike interactions. Such models have been developed for spike trains alone Truccolo et al. [2005]. Combining them with latent spectral dynamics would connect single-trial oscillatory fluctuations to population spiking patterns.

The biophysical relationship between spikes and fields involves volume conduction, synaptic currents, and network architecture that the current phenomenological model does not represent. Mechanistic models that derive field potentials from spiking activity—or vice versa—could provide more interpretable coupling parameters. For example, if the LFP reflects summed synaptic input to a local population, and spikes reflect output of that population, then the coupling coefficient *β*_*C*_ would have a concrete interpretation in terms of input-output gain. Forward models of the LFP based on multicompartment neuron simulations Einevoll et al. [2013] could be embedded within the state-space framework, though at substantially increased computational cost.

In summary, Joint SSMT provides a unified Bayesian framework for estimating time-resolved spectral dynamics and spike–field coupling from the same data stream. By combining multitaper LFP observations with spike timing through conditionally Gaussian augmentation, it yields frequency-specific coupling estimates with principled uncertainty quantification and improves spectrogram denoising at coupled bands. Across simulations and two primate datasets, Joint SSMT localizes coupling more sharply than PLV and SFC while retaining interpretability through its generative parameters.

## Acknowledgments

We thank Alexandra Bardon for helpful comments on the manuscript. This work was supported by the Office of Naval Research Multidisciplinary University Research Initiative (N00014-23-1-2768), the Army Research Office (W911NF2410228), the Freedom Together Foundation, and The Picower Institute for Learning and Memory. The propofol anesthesia data were collected as part of the study by Bastos et al. Bastos et al. [2021]. The paired-associates task data were collected as part of studies by Brincat and Miller Brincat and Miller [2015].

## A Notation

Three tables summarize the notation used throughout the paper. Table S1 lists indices, grids, and structural quantities. Table S2 defines the generative model variables. Table S3 covers inference, testing, and auxiliary quantities.

**Table S1:** Indices, time grids, and structural quantities.

| Symbol | Range / value | Meaning |
| --- | --- | --- |
| *Indices* |  |  |
| $n$ | $1, \dots, N_{\text{spk}}$ | Spike-bin index |
| $k$ | $1, \dots, K$ | Block (LFP window) index |
| $m$ | $1, \dots, M$ | Slepian taper index |
| $j$ | $1, \dots, J_f$ | Frequency-bin index |
| $h$ | $1, \dots, H$ | Spike-history lag index |
| $r$ | $1, \dots, R$ | Trial index |
| *Time grids* |  |  |
| $t$ | $\mathbb{R}_{\geq 0}$ | Continuous time (OU diffusion only) |
| $t_n = n \Delta_{\text{spk}}$ | — | Spike-bin center |
| $t_k = k \Delta_b$ | — | Block center |
| $\Delta_{\text{spk}}$ | e.g. 1–10 ms | Spike-bin width |
| $\Delta_b = J \Delta$ | e.g. 100–2000 ms | Block width ($J$ samples at interval $\Delta$) |
| *Dimensions and counts* |  |  |
| $N_{\text{spk}}$ | — | Total number of spike bins |
| $n_{\text{spk}}$ | — | Number of spikes ($\sum_n S_n$) |
| $K$ | — | Number of LFP blocks |
| $M$ | — | Number of Slepian tapers |
| $J_f$ | — | Number of frequency bins |
| $H$ | — | Number of spike-history lags |
| $R$ | — | Number of trials |

**Table S2:**
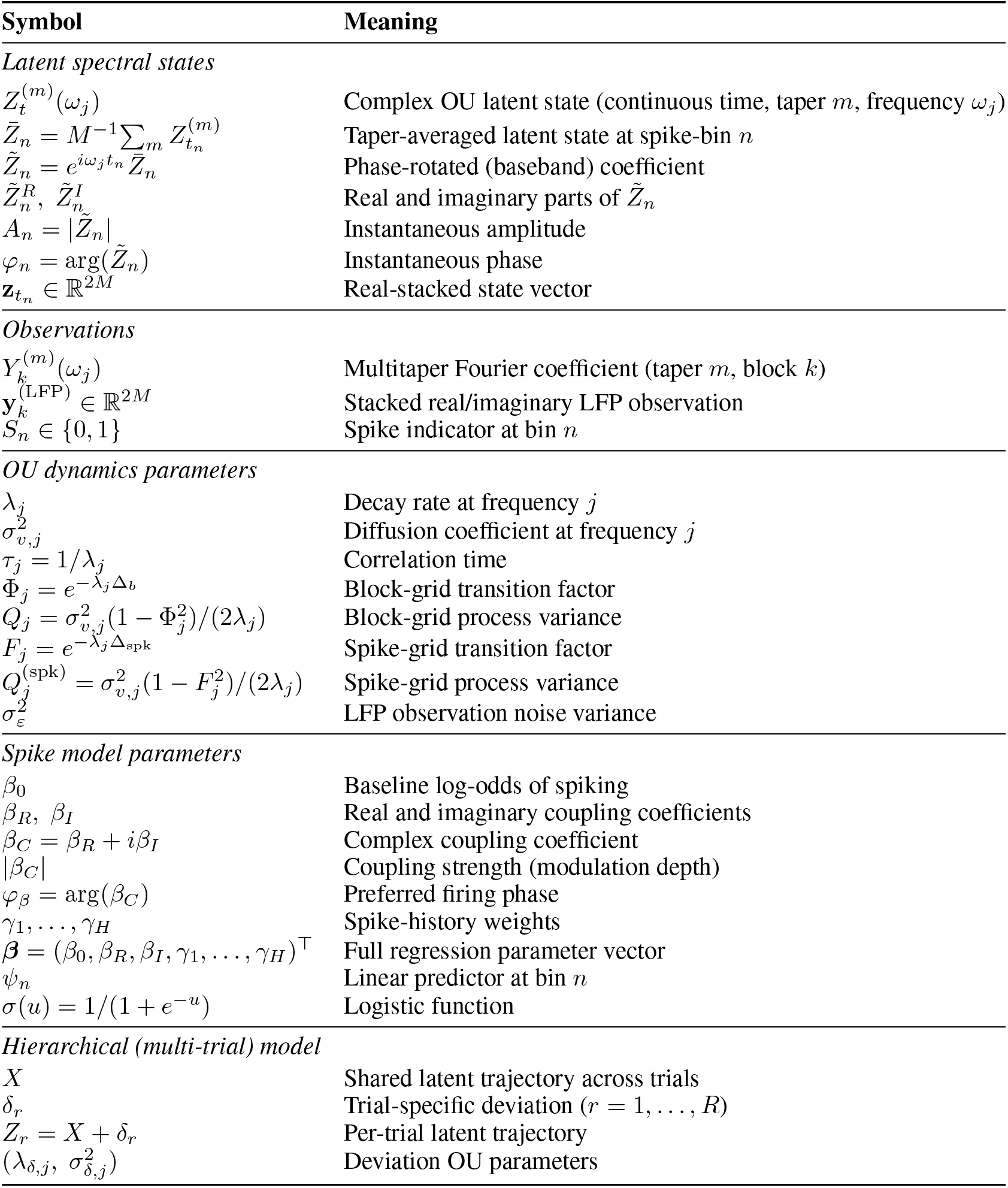
Generative model: latent states, observations, and parameters.

**Table S3:**
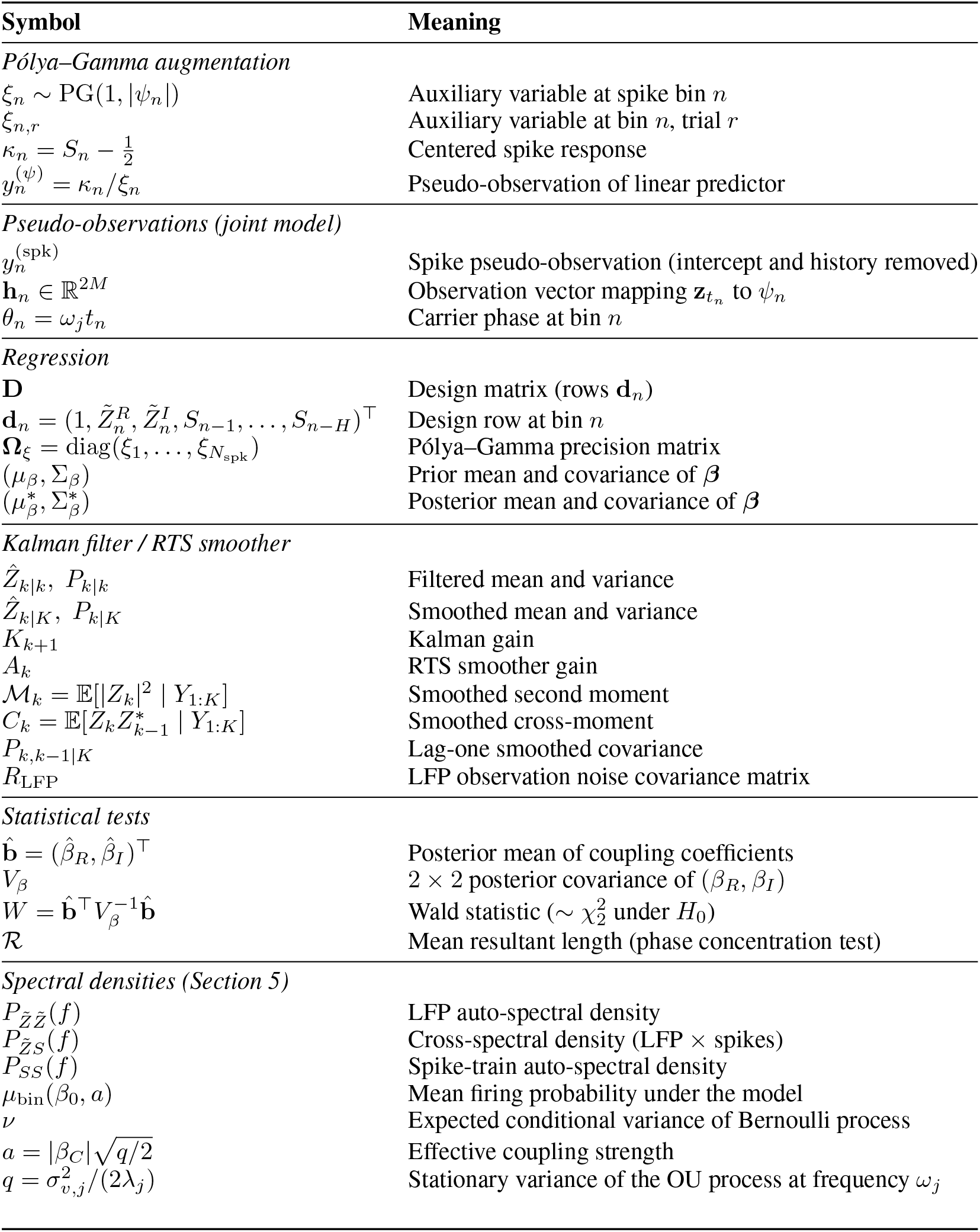
Inference quantities, augmentation variables, and statistical tests.

## B Inference details

This appendix derives the Kalman filtering, smoothing, and parameter estimation procedures. Section B.1 covers the LFP-only model. Section B.2 extends the framework to incorporate spike observations via Pólya–Gamma augmentation.

### B.1 LFP-only inference

We work with a single taper–frequency channel (*m, j*); because channels are conditionally independent, the results apply in parallel across all *M* × *J*_*f*_ channels. To simplify notation, we suppress the indices (*m, j*) and write *Z*_*k*_ for the latent state at block center *t*_*k*_, *Y*_*k*_ for the observed multitaper coefficient, Φ_*j*_ for the transition factor, *Q*_*j*_ for the process variance, and 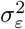 for the observation variance.

#### State-space model

The Ornstein–Uhlenbeck diffusion (3) discretized at the block spacing Δ_*b*_ yields the linear Gaussian state-space model

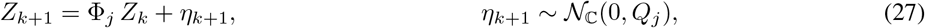

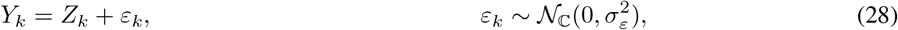

where *N*_ℂ_ (*µ, q*) denotes a proper complex normal distribution with mean *µ* and variance *q* = E[*X* − *µ* ^2^]. We derive the transition parameters by solving the OU SDE. For *λ*_*j*_ > 0, the solution starting from *Z*_*t*_ is

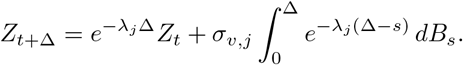

The stochastic integral is a zero-mean complex Gaussian with variance

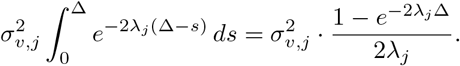

Setting Δ = Δ_*b*_ gives

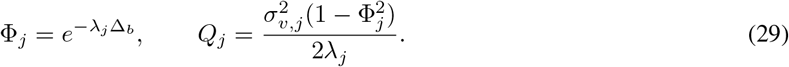

When *λ*_*j*_ = 0, the process is a complex Brownian motion with Φ_*j*_ = 1 and 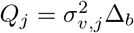.

#### Kalman filter

Let 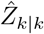 and *P*_*k*|*k*_ denote the filtered mean and variance of *Z*_*k*_ given observations *Y*_1:*k*_. We derive the Kalman recursions by computing conditional distributions.

Given 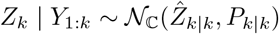 and the state equation (27), the one-step-ahead predictive distribution is

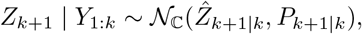

where

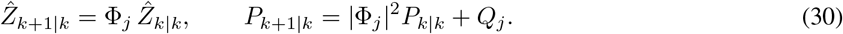

The first equality follows from linearity of expectation; the second from the independence of *η*_*k*+1_ and *Z*_*k*_ | *Y*_1:*k*_.

Given the predictive distribution and the observation *Y*_*k*+1_ = *Z*_*k*+1_ + *ε*_*k*+1_, the joint distribution of (*Z*_*k*+1_, *Y*_*k*+1_) given

*Y*_1:*k*_ is Gaussian. By standard conditioning formulas for complex Gaussians,

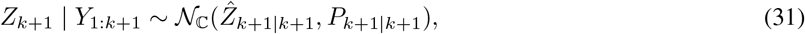

with

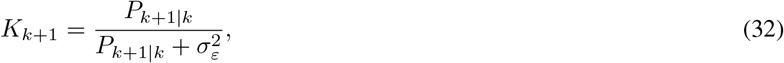

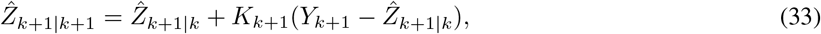

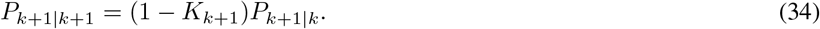

The Kalman gain *K*_*k*+1_ ∈ [0, 1] interpolates between the prediction and observation according to their relative precisions.

To verify (32)–(34), write the joint covariance of (*Z*_*k*+1_, *Y*_*k*+1_) given *Y*_1:*k*_:

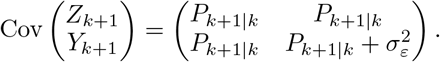

The conditional mean of *Z*_*k*+1_ given *Y*_*k*+1_ is 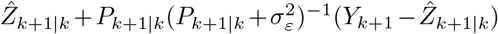, which gives (32)–(33). The conditional variance is 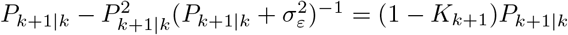, which gives (34).

We initialize the filter with a diffuse prior *Z*_0_ ~ *N*_ℂ_ (0, *P*_0|0_) where *P*_0|0_ is large, or with the stationary distribution of the OU process, 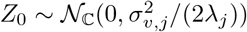 when *λ*_*j*_ > 0.

#### Rauch–Tung–Striebel smoother

The Kalman filter computes *p*(*Z*_*k*_ |*Y*_1:*k*_), but for parameter estimation we require the smoothed distribution *p*(*Z*_*k*_ | *Y*_1:*K*_) given all *K* observations. The Rauch–Tung–Striebel (RTS) algorithm computes these distributions via a backward recursion.

Let 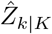 and *P*_*k*|*K*_ denote the smoothed mean and variance. The algorithm proceeds as follows. Initialize with 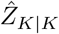 and *P*_*k*|*K*_ from the final filter step. For *k* = *K* − 1, …, 0, compute the smoothing gain

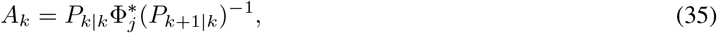

and update

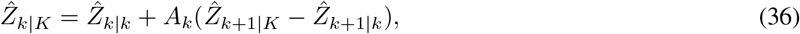

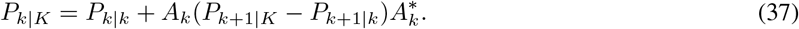

To derive (35)–(37), we use the Markov property and Bayes’ rule. The smoothed distribution satisfies

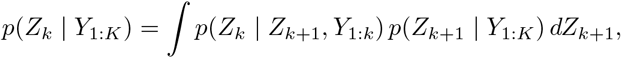

where the first factor uses the Markov property: given *Z*_*k*+1_, the state *Z*_*k*_ is conditionally independent of *Y*_*k*+1:*K*_. The conditional distribution *p*(*Z*_*k*_ | *Z*_*k*+1_, *Y*_1:*k*_) is obtained by inverting the state equation. From (27), the joint distribution of (*Z*_*k*_, *Z*_*k*+1_) given *Y*_1:*k*_ is

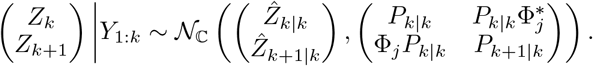

The off-diagonal term is Cov(*Z*_*k*_, *Z*_*k*+1_ | *Y*_1:*k*_) = Cov(*Z*_*k*_, Φ_*j*_*Z*_*k*_ + *η*_*k*+1_) = Φ_*j*_*P*_*k k*_, and its conjugate transpose is 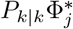. Conditioning on *Z*_*k*+1_ gives

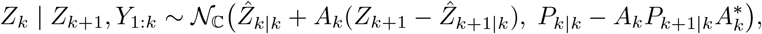

with *A*_*k*_ as in (35). The integral over the smoothed distribution 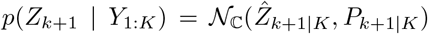 yields (36)–(37).

#### Forward filtering, backward sampling

For Gibbs sampling, we require draws from the joint smoothing distribution *p*(*Z*_0:*K*_ | *Y*_1:*K*_), not just the marginals. The forward filtering, backward sampling (FFBS) algorithm produces such draws efficiently [Carter and Kohn, 1994, Frühwirth-Schnatter, 1994].

Run the forward Kalman filter and store the filtered quantities 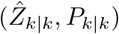 and predictive quantities 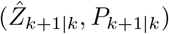 for all *k*. Then sample backward:

1. Draw 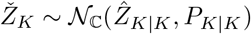
2. For *k* = *K* − 1, …, 0, draw

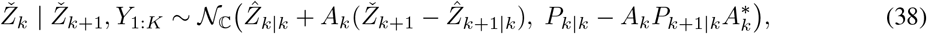

with *A*_*k*_ from (35).

The sample 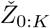 is an exact draw from the joint posterior *p*(*Z*_0:*K*_ | *Y*_1:*K*_).

#### EM updates

The EM algorithm requires the smoothed lag-one covariance *P*_*k,k* − 1 |*K*_ = Cov(*Z*_*k*_, *Z*_*k*−1_ |*Y*_1:*K*_). This quantity can be computed from the RTS backward pass. Using the law of total covariance and the backward conditional 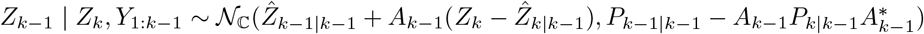, we obtain

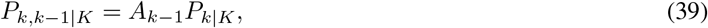

where *A*_*k*−1_ is real for the OU process.

Given smoothed moments, we update the model parameters 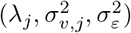 by maximizing the expected complete-data log-likelihood. Define the smoothed second moments

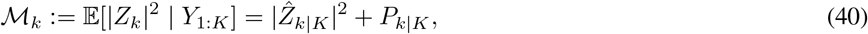

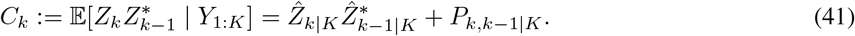

The complete-data log-likelihood for the state equation, summed over blocks *k* = 1, …, *K* and tapers *m* = 1, …, *M*, is

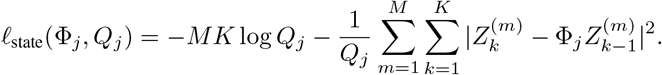

Taking expectations and differentiating with respect to Φ_*j*_ and *Q*_*j*_ yields the M-step updates. Setting ∂E[ℓ_state_]*/*∂Φ_*j*_ = 0 gives

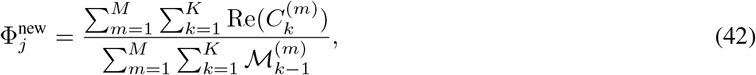

Setting ∂E[ℓ_state_]*/*∂*Q*_*j*_ = 0 gives

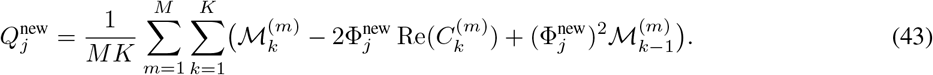

The continuous-time parameters are recovered by inverting (29):

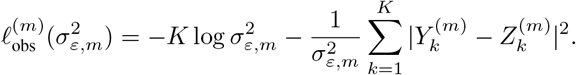

For the observation noise, the complete-data log-likelihood for taper *m* is

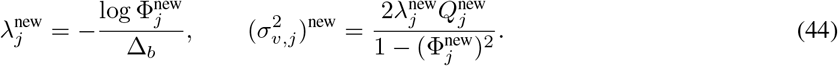

The M-step update is

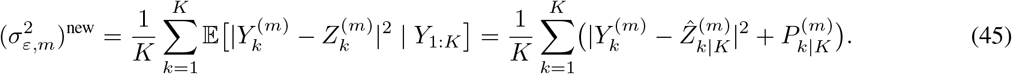

If the observation variance is shared across tapers, average (45) over *m*. When observation noise varies across tapers, replace 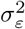 with 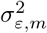 throughout.

### B.2 Joint inference with spike observations

When spike data are available, we work on the finer spike-bin grid {*t*_*n*_} with spacing Δ_spk_ ≪ Δ_*b*_. The Pólya–Gamma augmentation described in Section 2.2 converts each spike bin into a Gaussian pseudo-observation. Here we describe how these pseudo-observations enter the state-space framework.

#### State representation

To apply real-valued Kalman filtering, we represent the *M* complex taper-specific latent states at time *t*_*n*_ as a real vector by stacking real and imaginary parts:

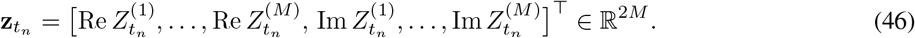

The OU dynamics discretized at Δ_spk_ become

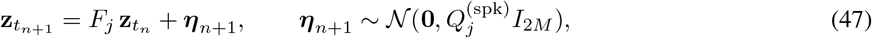

where 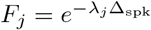 (a scalar applied elementwise) and 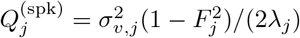. The tapers evolve independently with identical dynamics, so the transition matrix is *F*_*j*_*I*_2*M*_ and the process covariance is diagonal.

#### LFP observation rows

At block centers *t*_*k*_, the LFP provides a 2*M*-dimensional Gaussian observation. Stacking real and imaginary parts of the multitaper coefficients:

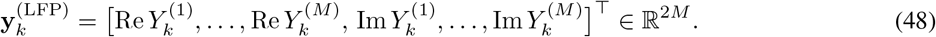

The observation model is

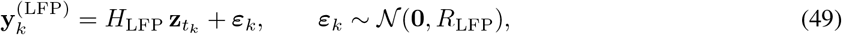

where *H*_LFP_ = *I*_2*M*_ and 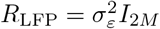 if the observation noise is shared across tapers. More generally, *R*_LFP_ can be diagonal with taper-specific variances.

#### Spike pseudo-observation rows

The Pólya–Gamma augmentation (Section 2.2) produces a Gaussian pseudo-observation at each spike bin. From (11), the pseudo-observation is

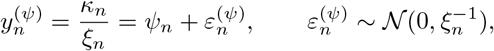

where 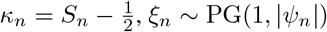, and the linear predictor is 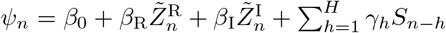.

To express *ψ*_*n*_ as a linear function of 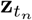, we derive the relationship between 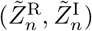 and the stacked state. Recall that 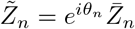 with *θ*_*n*_ = *ω*_*j*_*t*_*n*_ and 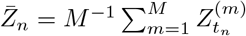. Writing 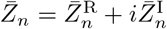, we have

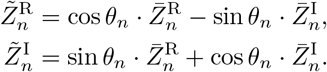

Since 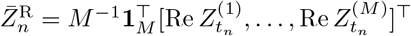 and similarly for 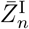, we can write

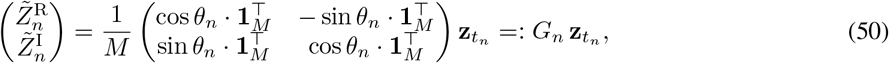

where *G*_*n*_ ∈ ℝ^2×2*M*^.

The linear predictor (excluding the known offset terms) is 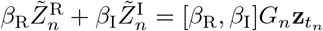. Defining

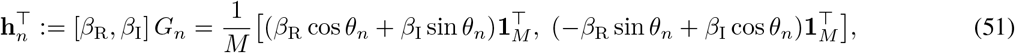

the spike pseudo-observation becomes

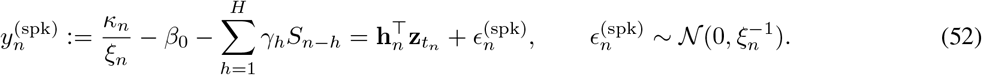

This is a scalar Gaussian observation with observation vector **h**_*n*_ and variance 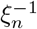.

#### Combined Kalman filter

On the spike-bin grid 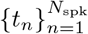, the latent state evolves according to (47). At each time *t*_*n*_, we apply:

1. *Prediction:* 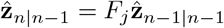 and 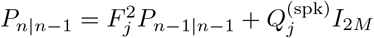.
2. *Spike update:* Apply the scalar observation (52) with Kalman gain

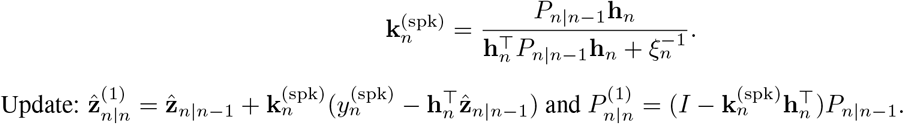

Update: 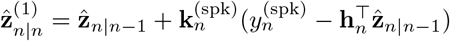 and 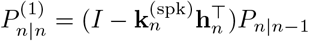.

- *LFP update (if t*_*n*_ *corresponds to a block center t*_*k*_): Apply the vector observation (49) with Kalman gain

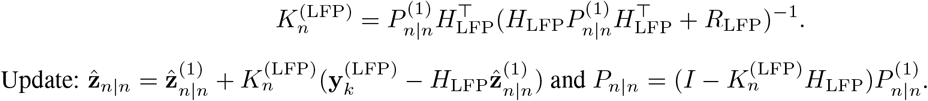

If no LFP observation is present at *t*_*n*_, set 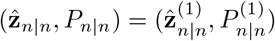. Because both observation types are conditionally Gaussian given the latent state, they can be applied in either order when they coincide; the final filtered distribution is the same.

The RTS smoother and FFBS algorithm extend directly to this multivariate setting. The smoothing gain becomes *A*_*n*_ = *P*_*n* |*n*_*F*_*j*_(*P*_*n*+1 |*n*_)^−1^ ℝ^2*M*×2*M*^, and the backward recursions (36)–(37) and (38) apply with vector/matrix operations.

The coupling coefficients ***β*** are updated via (17) in the main text, using either a sampled latent trajectory or the smoothed moments 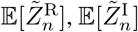, and their second moments derived from 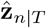 and *P*_*n*|*T*_ via the linear transformation (50).

## C Hierarchical trial inference

We implement the hierarchical model *Z*_*r*_ = *X* + *δ*_*r*_ via a two-pass approximate decomposition. The first pass pools observations across trials to estimate the shared process *X*; the second pass estimates each trial-specific trajectory *Z*_*r*_ independently, from which we recover the deviation *δ*_*r*_ = *Z*_*r*_ − *X*. Both passes operate on the uniform spike-bin grid *{t*_*n*_*}*, with LFP block centers *{t*_*k*_*}* treated as a subset of this grid. The OU hyperparameters 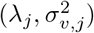 for the shared process and 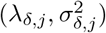 for the deviations are estimated via hierarchical EM prior to the iterative procedure and held fixed thereafter.

### C.1 Two-pass trajectory estimation

To estimate the shared trajectory, we pool observations across trials using precision weighting. For LFP, define weights 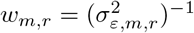 and form pooled observations

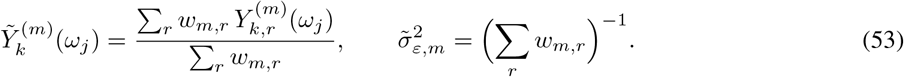

For spikes, draw Pólya–Gamma auxiliary variables *ξ*_*n,r*_ ~ PG(1, *ψ*_*n,r*_) using current parameters. Define the spike pseudo-observation for each trial,

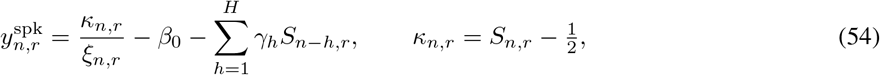

and pool across trials:

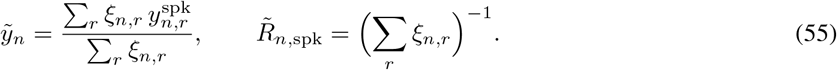

Run a joint Kalman filter and RTS smoother on the pooled observations 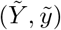 with OU dynamics 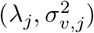 to obtain the smoothed shared trajectory 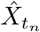 and its posterior covariance.

For each trial *r* separately, run a Kalman filter and RTS smoother using the per-trial LFP observations 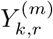 and spike pseudo-observations 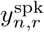. We use combined OU dynamics that capture both shared and trial-specific variability, with *λ*_*Z,j*_ = min(*λ*_*j*_, *λ*_*δ,j*_) and 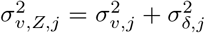. This yields the smoothed trial trajectory 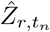 and its posterior covariance.

Recover the trial deviation as

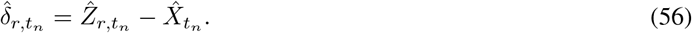

The posterior variance of *δ*_*r*_ is approximated conservatively as 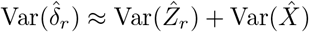.

### C.2 Parameter updates

#### Coupling coefficients

Given smoothed trajectories, we update the coupling coefficients ***β*** = (*β*_0_, *β*_R_, *β*_I_)^⊤^ using the posterior moments of the latent predictors. For each trial, form the taper-averaged, phase-rotated predictor

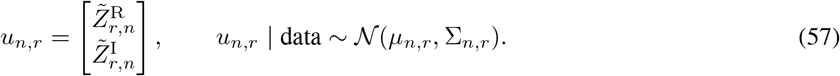

With design vector 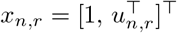 and prior ***β*** ~ *N* (*µ*_0_, ∑_0_), compute the posterior precision and information:

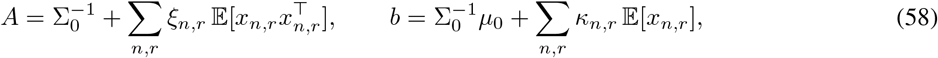

where the required expectations are

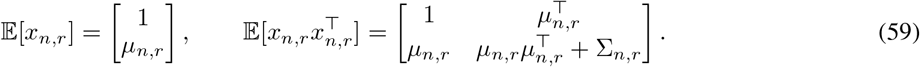

The posterior mean is 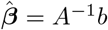. Including ∑_*n,r*_ in the second-moment computation propagates latent uncertainty into the coupling estimate.

### C.3 Algorithm summary

One outer iteration proceeds as follows:

i. Draw *ξ*_*n,r*_ ~ PG(1, *ψ*_*n,r*_) for all (*n, r*) using current ***β*** and predictors 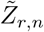.
ii. *Pass 1:* Pool observations via (53)–(55); run Kalman smoother to obtain 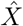.
iii. *Pass 2:* For each trial *r*, run Kalman smoother to obtain 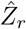; recover 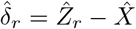.
iv. Update ***β*** via (58)–(59).

Iterate until convergence or for a fixed number of outer iterations.

## D Supplementary methods: single-trial analysis

This appendix describes the simulation protocol, inference configuration, and baseline methods for the single-trial analyses in Section 3.

### D.1 Simulation protocol

We generated synthetic data to evaluate the joint inference framework under controlled conditions where ground-truth coupling parameters are known.

#### LFP generation

The latent spectral state at each frequency evolves according to the Ornstein–Uhlenbeck diffusion (3). We simulated *J* = 6 signal frequencies: *{*7, 11, 19, 27, 35, 43*}* Hz. Of these, four are designated as coupled (*{*11, 19, 27, 43*}* Hz) and two are designated as uncoupled (*{*7, 35*}* Hz). The OU parameters are *λ*_*j*_ = *π* · 0.03 ≈ 0.094 rad/s, corresponding to a half-bandwidth of 0.03 Hz, and *σ*_*v,j*_ = 3.0 for all signal bands.

At each time *t*_ℓ_ = ℓΔ with Δ = 1 ms, we synthesized the observed LFP as

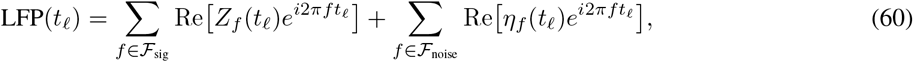

where ℱ_sig_ = *{*7, 11, 19, 27, 35, 43*}* Hz and ℱ_noise_ = *{*1, 2, …, 60*} \* ℱ_sig_. For signal frequencies, we added complex Gaussian observation noise with standard deviation *σ*_ε_ = 15. For noise-only frequencies, the state is 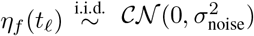 with *σ*_noise_ = 30. This construction ensures that the observed LFP contains oscillatory structure at signal frequencies embedded in broadband noise.

#### Spike generation

We simulated *N*_*u*_ = 5 units. Each unit couples to *k*_active_ = 3 frequencies drawn without replacement from the four coupled bands. The coupling coefficient for unit *s* at frequency *f* is

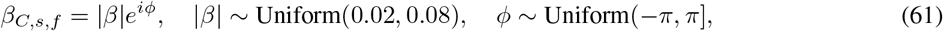

for coupled pairs, and *β*_*C,s,f*_ = 0 otherwise.

Spikes were generated in 1 ms bins according to the Bernoulli model (6). The baseline log-odds was *β*_0_ = −3.5, yielding mean firing rates of approximately 3–6 Hz depending on the coupling-induced modulation. We included a spike-history filter with *H* = 20 lags and an exponentially decaying inhibitory kernel,

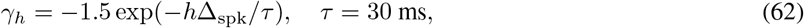

which enforces a relative refractory period. The total simulation duration was *T* = 300 s.

### D.2 Inference configuration

#### Multitaper settings

We analyzed the LFP using a frequency grid from 1 to 60 Hz in 1 Hz steps. The multitaper window was Δ_*b*_ = 2 s with time-bandwidth product *NW* = 2, yielding *M* = 3 Slepian tapers and a spectral resolution of 2 Hz. Windows were non-overlapping, producing *K* = 150 time blocks.

#### CT-SSMT inference

For the LFP-only model, we ran the EM algorithm to estimate the OU parameters 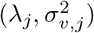 and observation noise 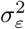 at each frequency independently. The Kalman smoother then computed posterior means and variances of the latent trajectory at block centers.

#### Joint SSMT inference

For the combined spike–LFP model, we alternated between three steps: (i) Pólya–Gamma sampling, drawing *ξ*_*n*_ ∼ PG(1, |*ψ*_*n*_) for each spike bin using the method of Polson et al. [2013]; (ii) coupling update, sampling ***β*** from its Gaussian conditional posterior (17) with prior 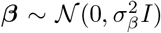 where *σ*_*β*_ = 1; and (iii) latent trajectory update, running the Kalman smoother on the spike-bin grid with both LFP and spike pseudo-observations. We performed 1000 iterations of this scheme, discarding the first 60% as burn-in. Point estimates are posterior means computed from the retained samples. Credible intervals are sample quantiles.

### D.3 Baseline methods

#### Phase-locking value (PLV)

For each unit–frequency pair, we bandpass filtered the LFP using a second-order Butterworth filter with bandwidth equal to 2 · *NW/*Δ_*b*_ = 2 Hz centered at the frequency of interest. We extracted instantaneous phase via the Hilbert transform and computed

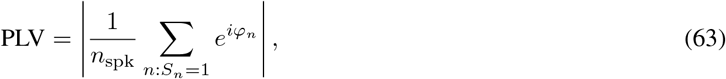

where *φ*_*n*_ is the LFP phase at spike time *n*.

#### Spike-field coherence (SFC)

We computed coherence between the spike train, treated as a binary time series at 1 ms resolution, and the LFP using Welch’s method with 2-second segments and 50% overlap. Coherence at frequency *f* is

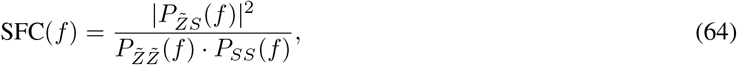

where 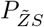 is the cross-spectral density and 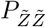, *P*_*SS*_ are the auto-spectral densities.

### D.4 Statistical testing

#### Joint model: Wald test

Let 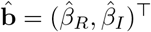 denote the posterior mean vector and let *V*_*β*_ denote the 2 × 2 posterior covariance matrix estimated from the MCMC samples. The Wald statistic is

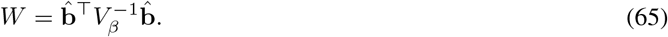

This quantity has a direct Bayesian interpretation: for a bivariate Gaussian posterior, the *α*-credible ellipse is the set 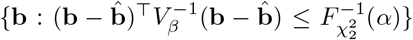. The origin lies inside this ellipse if and only if 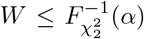. Thus 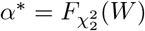 is the smallest credible level whose ellipse contains the origin, and we report *p* = 1 − *α*^∗^. A small ridge (10^−10^ · **I**_2_) is added to *V*_*β*_ for numerical stability.

#### Joint model: phase concentration test

The Wald test assesses whether the posterior mean deviates from zero but does not directly measure phase consistency across posterior samples. We therefore also computed a phase concentration statistic using the Rayleigh test. Let 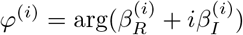 denote the phase of the coupling coefficient at MCMC iteration *i*. The mean resultant length is

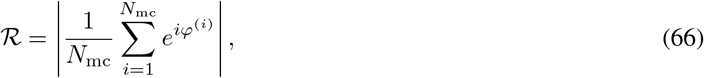

where *N*_mc_ is the number of posterior samples. Values of ℛ near 1 indicate consistent preferred phase across posterior samples, while values near 0 indicate dispersed phases. Under the null hypothesis that the posterior phases are uniformly distributed, 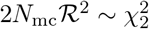, yielding *p* = exp(−*N*_mc_ℛ^2^).

#### PLV: Rayleigh test

The Rayleigh test assesses uniformity of the circular distribution of spike phases. Given *n*_spk_ spikes with LFP phases 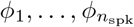 at spike times, the PLV equals the mean resultant length 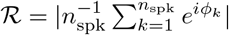. Under the null hypothesis of uniform spike phases, 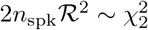, yielding *p* = exp(−*n*_spk_ · PLV^2^).

#### SFC: parametric test

For coherence estimated from *L* segments, the quantity *L*·SFC*/*(1 SFC) follows an *F*_2,2*L*2_ distribution under the null hypothesis of zero coherence.

#### Permutation tests

We also computed non-parametric *p*-values for PLV and SFC using two surrogate procedures. In the circular shift procedure, we circularly shifted the spike train by a random offset drawn uniformly from [*T/*4, 3*T/*4] and recomputed the statistic; this was repeated 500 times. In the local jitter procedure, we added independent uniform jitter drawn from [−25 ms, 25 ms] to each spike time and recomputed the statistic; this was also repeated 500 times. The *p*-value is (1 + #{surrogate ≥ observed})*/*(1 + *n*_perm_).

### D.5 Performance metrics

#### Spectrogram correlation

To quantify how well each method tracks the true spectral dynamics, we computed Pearson correlations between estimated and ground-truth power in non-overlapping time windows of duration Δ_corr_ = 20 s. For each signal frequency *j* and window *w*, let 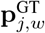 and 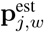 denote the vectors of ground-truth and estimated power values at the fine temporal resolution (1 ms). The correlation is

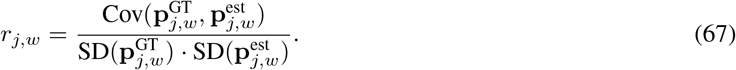

We report boxplots of *{r*_*j,w*_*}*_*w*_ for each frequency and method.

#### Spectrogram visualization

All spectrograms display power in decibels, 10 log_10_(|*A*_*j,k*_|^2^), where *A*_*j,k*_ is the amplitude at frequency *j* and time block *k*. For ground-truth simulations, *A*_*j,k*_ = |*Z*_*j*_(*t*_*k*_)| is the true latent amplitude. For multitaper estimates, 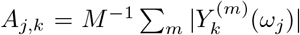. For CT-SSMT estimates, 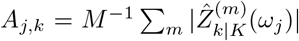 where 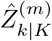 is the posterior mean from the Kalman smoother. Color scales are set independently for each method to span the 0th to 100th percentile of that method’s power values, ensuring visual comparability.

### D.6 Propofol anesthesia data analysis

#### Recording and experimental protocol

We analyzed multielectrode recordings from macaque cortex during propofol-induced loss of consciousness, a dataset previously analyzed in Bastos et al. [2021]. LFP and single-unit activity were recorded from ventrolateral prefrontal cortex (vlPFC) and posterior parietal area 7b using chronically implanted electrode arrays. The animal received intravenous propofol infusion while neural activity was continuously monitored. LFP signals were sampled at 1000 Hz. Surgical, pharmacological, and experimental details appear in Bastos et al. [2021].

#### Epoch definitions

The recording was partitioned into six epochs of approximately 600 s each, defined by drug administration timing and behavioral state markers. Baseline comprised the period before drug infusion, approximately 600–1110 s from recording onset. Drug Onset spanned approximately 1120–1720 s, immediately following infusion start. LOC 1, approximately 1720–2320 s, encompassed behavioral loss of consciousness as indicated by eye closure. LOC 2, approximately 4120–4720 s, represented deep anesthesia in the 600 s preceding drug cessation. Drug Stop, approximately 4720–5320 s, covered the period immediately following infusion termination. Recovery, approximately 5320–5920 s, captured the return to consciousness as indicated by eye opening.

#### Unit selection

We included units with mean firing rates of at least 2 Hz computed over the full recording duration. This criterion was applied globally rather than epoch-by-epoch to ensure consistent unit identity across conditions; a unit included in any epoch was retained for analysis in all epochs. This procedure yielded *N*_*u*_ = 13 units for the analyses presented in Section 3.

#### Spectral and inference parameters

The frequency grid ranged from 0.4 to 30 Hz in 0.4 Hz steps to capture the slow oscillations characteristic of propofol anesthesia. Multitaper windows of 2 s duration with time-bandwidth product *NW* = 2 yielded *M* = 3 Slepian tapers and a spectral resolution of 2 Hz. For the joint model, we ran 1500 MCMC iterations with 60% burn-in and 5 Kalman filter refreshes during sampling.

#### Baseline methods

PLV and SFC were computed as described in Section D.3, with filter bandwidths matched to the spectral resolution. The Rayleigh test (PLV) and coherence *F*-test (SFC) provided parametric *p*-values. Permutation tests used 500 circular shifts per unit–frequency pair, with shift offsets drawn uniformly from the middle half of each epoch duration to avoid edge effects.

## E Supplementary methods: trial-structured data analysis

This appendix describes the simulation protocol and analysis configuration for the trial-structured experiments in Section 4.

### E.1 Trial-structured simulation protocol

We generated synthetic data to evaluate the hierarchical inference framework under conditions where the shared and trial-specific components are known.

#### LFP generation

The latent spectral state at each frequency decomposes into shared and trial-specific components according to the hierarchical model (18). We simulated six signal frequencies partitioned into two groups: four couplable bands at 11, 19, 27, and 43 Hz that can drive spike timing, and two signal-only bands at 7 and 35 Hz that carry true oscillatory signal but never couple to spikes (*β*_*C*_ = 0). This design tests whether the inference correctly identifies coupling only where it exists, without false positives at signal-only bands.

The shared process *X*_*j*_(*t*) evolves according to the OU diffusion (3) with half-bandwidth Δ*f*_*j*_ = 0.05 Hz, corresponding to *λ*_*j*_ = *π* ·0.05 ≈ 0.157 rad/s, and diffusion coefficient *σ*_*v,j*_ = 4.0 for all signal bands. The trial-specific deviations *δ*_*r,j*_(*t*) evolve with fourfold faster decay (*λ*_*δ,j*_ = 4*λ*_*j*_) and diffusion coefficient *σ*_*δ,j*_ = 5.0. At each time *t*_ℓ_ = ℓΔ with Δ = 1 ms, we synthesized the observed LFP as

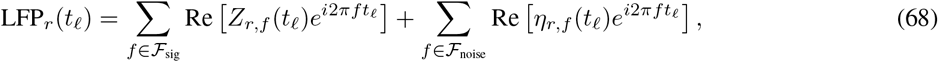

where ℱ_sig_ = *{*7, 11, 19, 27, 35, 43*}* Hz contains the signal bands and ℱ_noise_ = *{*1, 2, …, 60*} \* ℱ_sig_ contains noise-only bands with 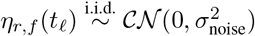 and *σ*_noise_ = 10. Complex Gaussian observation noise with *σ*_*ε*_ = 5 was added to signal bands.

#### Spike generation

We simulated *N*_*u*_ = 5 units, each coupling to *k*_active_ = 3 frequencies drawn without replacement from the four couplable bands. For unit *s* at coupled frequency *f*, the coupling coefficient is *β*_*C,s,f*_ = |*β*|*e*^*iϕ*^ with magnitude |*β*| ~ Uniform(0.02, 0.15) and phase *ϕ* ~ Uniform(−*π, π*]; uncoupled pairs have *β*_*C,s,f*_ = 0.

Spikes were generated in 1 ms bins according to the Bernoulli model (6) with baseline log-odds *β*_0_ ~ N (−2.0, 0.4^2^), yielding mean firing rates of approximately 3–8 Hz depending on coupling-induced modulation. We included a spike-history filter with *H* = 20 lags and an exponentially decaying inhibitory kernel *γ*_*h*_ = − 1.5 exp(−*h*Δ_spk_*/τ*) with *τ* = 30 ms, which enforces a relative refractory period. The total simulation comprised *R* = 100 trials of *T* = 10 s each.

#### Inference configuration

We analyzed the LFP using a frequency grid from 1 to 61 Hz in 2 Hz steps (*J*_*f*_ = 30 bands). The multitaper window was Δ_*b*_ = 0.4 s with time-bandwidth product *NW* = 1, yielding *M* = 1 Slepian taper and a spectral resolution of 2.5 Hz. For the hierarchical joint model, we ran EM iterations to estimate the OU parameters for both shared and trial-specific components, followed by MCMC sampling with 1000 warmup iterations, 5 refresh cycles of 200 iterations each, trace thinning by a factor of 2, and burn-in fraction of 0.7. Posterior means and 95% credible intervals were computed from the retained samples.

#### Baseline methods

We computed PLV and SFC using the same procedures as for the single-trial analysis (Appendix D), pooling spike phases and coherence estimates across trials. The Rayleigh test (PLV) and coherence F-test (SFC) provided parametric *p*-values. We also computed permutation-based *p*-values using circular shifting of the spike train relative to the LFP within each trial, with 500 permutations per unit–frequency pair.

### E.2 SSPA associative learning task analysis

#### Task and recording

In the sample-sample paired associates (SSPA) task Brincat and Miller [2015], macaques learned arbitrary associations between pairs of visual stimuli. On each trial, a sample stimulus appeared, followed by a delay period, and then a test stimulus. The animal indicated via saccade whether the test stimulus matched the learned associate of the sample. Multielectrode recordings were obtained from electrode arrays in hippocampal subfields (CA3), subiculum (Sub), the head of the caudate nucleus (hCd), the tail of the caudate (tCd), and ventral prefrontal cortex (PFCv). We used one session from the recording as a demonstration. LFP signals were sampled at 1000 Hz.

#### Trial and unit selection

We selected the last 400 valid trials classified as SSPA task, novel (not previously over-trained), and not flagged as bad. All behavioral conditions (match/nonmatch, correct/incorrect) were combined to maximize statistical power for coupling detection. The analysis window spanned −0.5 to 3.5 s relative to sample onset, capturing pre-stimulus baseline, stimulus presentation, delay, and response epochs. For each electrode, we included units with mean firing rates of at least 1 Hz computed over this window and pooled across selected trials.

#### Spectral and inference parameters

The frequency grid ranged from 1 to 80 Hz in 2 Hz steps (*J*_*f*_ = 40 bands). Multitaper windows of 300 ms duration with time-bandwidth product *NW* = 1 provided a spectral resolution of approximately 3.3 Hz. We ran 200 EM iterations for the CT-SSMT parameters, followed by MCMC with 1000 warmup iterations, 5 refresh cycles of 200 inner steps each, trace thinning by a factor of 2, and burn-in fraction of 0.5. ARD-style shrinkage priors were applied with *a*_0_ = *b*_0_ = 0.01, and Wald selection used *α* = 0.1 for feature screening. Results were saved electrode-by-electrode, with coupling parameters, spectral dynamics, and metadata stored separately for downstream analysis.

#### Baseline methods

PLV and SFC were computed as described in Section D. The Rayleigh test provided parametric *p*-values for PLV, and the coherence F-test for SFC. Permutation tests used trial shuffling. Spike trains were randomly reassigned to different trials while LFP signals retained their original trial labels, breaking the spike-LFP correspondence while preserving within-trial temporal structure. We used 500 permutations per unit–frequency pair.

## F Proofs for section 5

### F.1 Phase-locking values

#### Proof of Proposition 1

Since *Z* ~ *CN* (0, *q*) is circularly symmetric, we may write 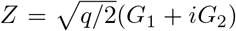 where 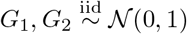. We rotate coordinates to align with the coupling direction. Assuming *β*_‖_ = |*β* | > 0, define

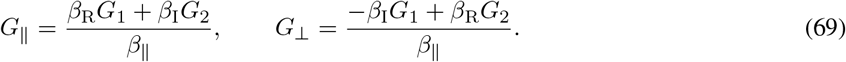

This is an orthogonal transformation, so (*G*_‖_, *G*_⊥_) remains a pair of independent standard normals. In these coordinates, the linear predictor simplifies:

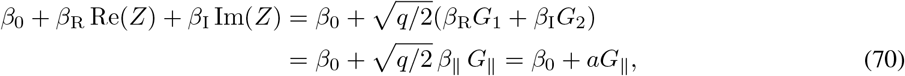

where 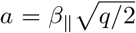. Thus the spike probability *σ*(*β*_0_ + *aG*_‖_) depends only on the component *G*_‖_ aligned with the coupling vector.

Inverting the rotation gives *G*_1_ + *iG*_2_ = (*β*_*C*_*/β*_∥_)(*G*_∥_ + *iG*_⊥_), so the unit phase factor becomes

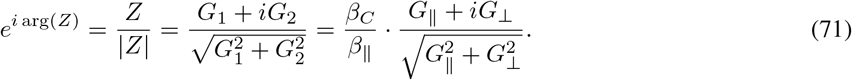

Using E[*S* | *Z*] = *σ*(*β*_0_ + *aG*_∥_), the numerator of the PLV is

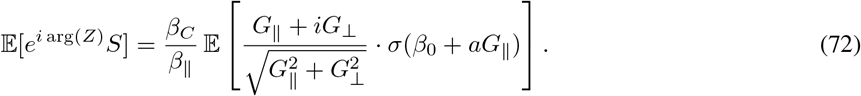

We evaluate the real and imaginary parts separately. For the imaginary part, we condition on *G*_∥_ = *g* and note that 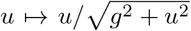 is an odd function. Since *G*_⊥_ ~ *N* (0, 1) is symmetric about zero, this conditional expectation vanishes, and hence the imaginary part is zero.

For the real part, we apply the tower property E[*X*] = E[E[*X* | *G*_∥_]]:

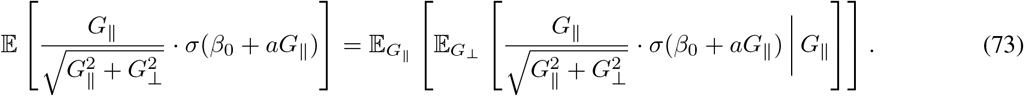

Conditioning on *G*_∥_ = *g* treats both *g* and *σ*(*β*_0_ + *ag*) as constants, so they factor out of the inner expectation:

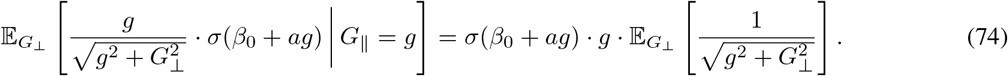

By independence, the conditional distribution of *G*_⊥_ given *G*_∥_ is just *N* (0, 1), so the inner expectation equals *h*(*g*) = E_*U*_ [(*g*^2^ + *U* ^2^)^−1*/*2^]. Taking the outer expectation over *G*_∥_:

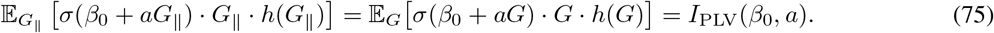

The denominator of the PLV is the mean spike probability:

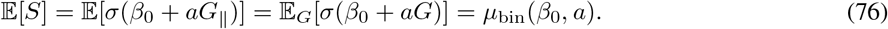

Combining these results,

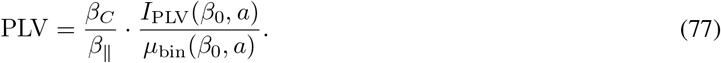

Since *I*_PLV_(*β*_0_, *a*) and *µ*_bin_(*β*_0_, *a*) are both real and positive, the phase of the PLV equals arg(*β*_*C*_*/β*_∥_) = arg(*β*_*C*_).

#### Remark (Closed form for *h*(*g*))

The function *h*(*g*) = E_*U*_ [(*g*^2^ + *U* ^2^)^−1*/*2^] admits the closed form

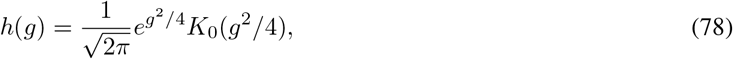

where *K*_0_ is the modified Bessel function of the second kind, defined by 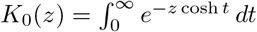. The identity follows from the substitution *u* = *g* sinh(*t/*2) in the defining integral for *h*(*g*). In practice, the expectations *µ*_bin_(*β*_0_, *a*) and *I*_PLV_(*β*_0_, *a*) are computed via Gauss–Hermite quadrature.

##### Proof of Corollary 1

The baseband OU process (3) has stationary distribution 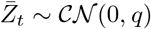 with 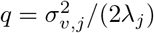. The phase-rotated process 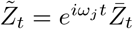 is obtained by multiplying by a deterministic unit-modulus factor. Since the complex Gaussian distribution *CN* (0, *q*) is circularly symmetric, we have 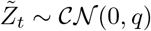 for all *t*, and proposition 1 applies with 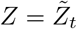.

The OU process is geometrically ergodic with unique stationary distribution *CN* (0, *q*). Since the spike-conditional expectation 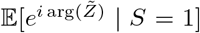 is a bounded function of 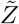, and 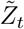 inherits ergodicity from 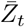, the empirical PLV converges almost surely to 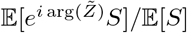 as the observation period grows.

Finally, the PLV magnitude depends on (*β*_0_, *β*_∥_, *q*) through the combination 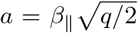, and the OU parameters (*λ*_*j*_, *σ*_*v,j*_) enter only through 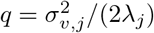. The frequency *ω*_*j*_ determines the phase rotation but does not affect *q* or any quantity in (20), so different frequencies with the same *q* yield identical PLV magnitudes.

Now this section

### F.2 Spike-field coherence

We work with the discrete-time model where 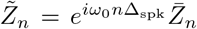 is the phase-rotated OU process with stationary variance 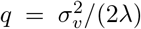, and spikes are generated according to 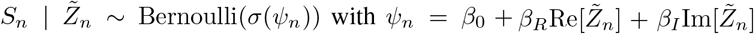. Define the complex coupling coefficient *β*_*C*_ = *β*_*R*_ + *iβ*_*I*_ and set 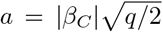. The key quantities are the mean firing probability *µ*_bin_ = E[*σ*(*β*_0_ + *aG*)] and the expected conditional variance ν = E[*σ*(*β*_0_ + *aG*)(1 − *σ*(*β*_0_ + *aG*))] where *G* ~ *N* (0, 1).

#### Proof of Proposition 2

We compute the cross-covariance 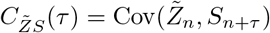 and take its Fourier transform. By the law of total covariance,

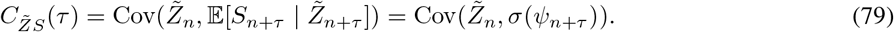

To evaluate this, we apply Stein’s lemma: for jointly Gaussian (*X, Y*) and differentiable *g*,

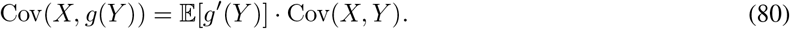

In our setting 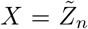 and *Y* = *ψ*_*n*+*τ*_, which are jointly Gaussian since *ψ*_*n*+*τ*_ is a linear function of the Gaussian process 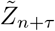. We need 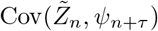.

Writing 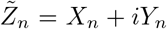 where 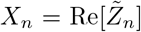 and 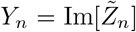, the linear predictor is *ψ*_*n*_ = *β*_0_ + *β*_*R*_*X*_*n*_ + *β*_*I*_*Y*_*n*_. For a proper complex Gaussian, Var(*X*_*n*_) = Var(*Y*_*n*_) = *q/*2 and Cov(*X*_*n*_, *Y*_*n*_) = 0. The autocovariance of 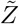 is 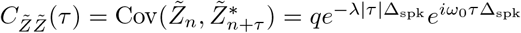, so

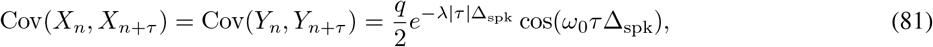

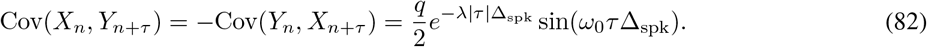

Computing the covariance with the linear predictor:

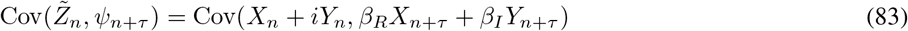

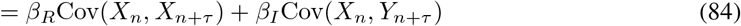

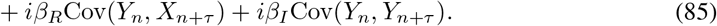

Substituting and simplifying using Euler’s formula:

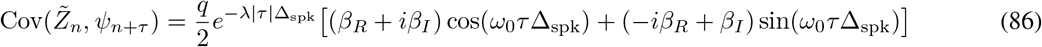

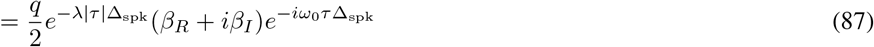

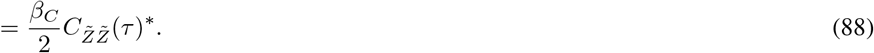

Since *σ*^*′*^(*x*) = *σ*(*x*)(1 − *σ*(*x*)), we have E[*σ*^*′*^(*ψ*_*n*_)] = E[*σ*(*ψ*_*n*_)(1 − *σ*(*ψ*_*n*_))] = ν. Applying Stein’s lemma:

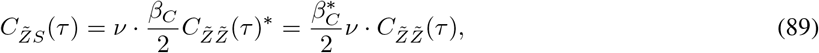

where the last equality uses the conjugate symmetry 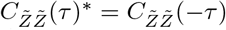 and that we can absorb the conjugate into *β*_*C*_. Taking the Fourier transform, the cross-spectral density is

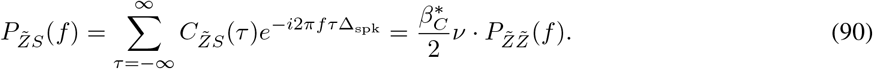

#### Proof of Proposition 3

Decompose the spike train as *S*_*n*_ = *λ*_*n*_ + *ϵ*_*n*_ where *λ*_*n*_ = *σ*(*ψ*_*n*_) is the instantaneous firing rate and *ϵ*_*n*_ = *S*_*n*_ − *λ*_*n*_ is the Bernoulli residual. The spike spectrum decomposes as *P*_*SS*_(*f*) = *P*_*λλ*_(*f*) + *P*_*ϵϵ*_(*f*) since Cov(*λ*_*n*_, *ϵ*_*n*+*τ*_) = 0 for all *τ*.

For the Bernoulli noise spectrum, the residual satisfies 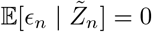 and 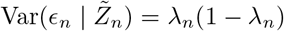. Since *ϵ*_*n*_ is conditionally independent across time given 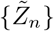, we have *C*_*ϵϵ*_(*τ*) = 0 for *τ* ≠ 0 and *C*_*ϵϵ*_(0) = E[*λ*_*n*_(1 − *λ*_*n*_)] = ν.

The power spectral density is therefore white:

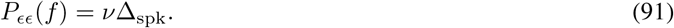

For the rate spectrum, the autocovariance is 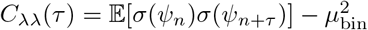. The inputs (*ψ*_*n*_, *ψ*_*n*+*τ*_) are jointly Gaussian with common variance 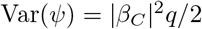 and normalized correlation

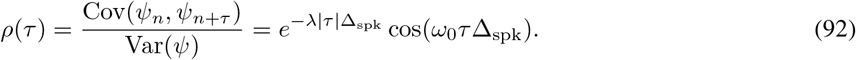

Price’s theorem states that for jointly Gaussian (*U, V*) with unit variance and correlation *ρ*,

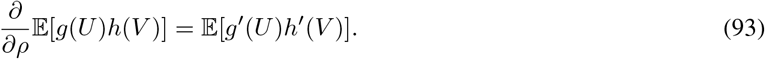

Applying this to standardized inputs with *g* = *h* = *σ* and integrating from *ρ* = 0:

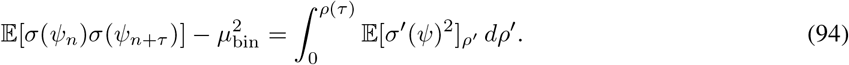

To leading order in *ρ*, the integrand evaluated at *ρ*^*′*^ = 0 gives E[*σ*^*′*^(*ψ*)]^2^ = ν^2^, yielding

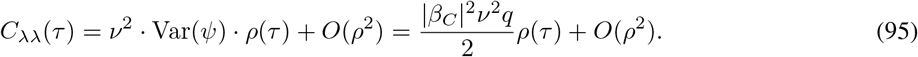

The correlation 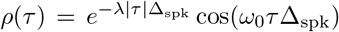 has a Lorentzian spectrum centered at ±*f*_0_ with *P*_*ρ*_(*f*_0_) = 1*/λ*. Therefore

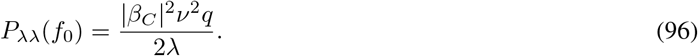

Combining the two components:

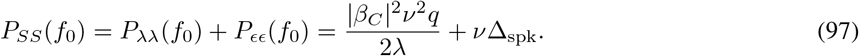

#### Proof of proposition 4

From Proposition 2, the cross-spectrum magnitude squared is

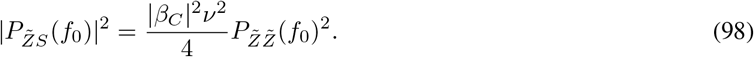

Substituting into the coherence definition with 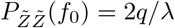:

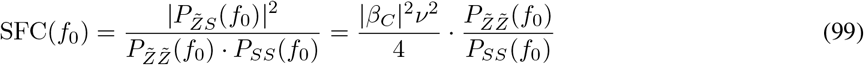

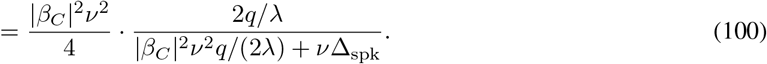

Let *A* = |*β*_*C*_|^2^ν^2^*q/*(2*λ*) denote the rate modulation power. Then

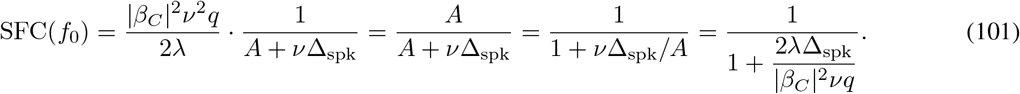

#### Remark

The closed-form SFC depends on (*β*_0_,| *β*_*C*_|, *q, λ*, Δ_spk_) but not on the carrier frequency *ω*_0_. This frequency independence holds because all spectral quantities at *f*_0_—namely 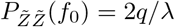, the cross-spectrum, and the spike spectrum—depend only on the baseband OU parameters. Changing the carrier shifts where the spectral peak occurs but does not change its height. The limiting behavior confirms physical intuition. As |*β*_*C*_| → ∞, the coupling becomes perfect and SFC → 1. As |*β*_*C*_| → 0, there is no coupling and SFC → 0. As *λ* → 0, the OU process becomes slow with a sharp spectral peak, concentrating signal power at *f*_0_ and yielding SFC → 1. Conversely, as *λ* → ∞, the spectrum flattens, signal power spreads across frequencies, and SFC → 0.

## G Supplementary figures

### G.1 Propofol induced anesthesia

**Figure S1:**
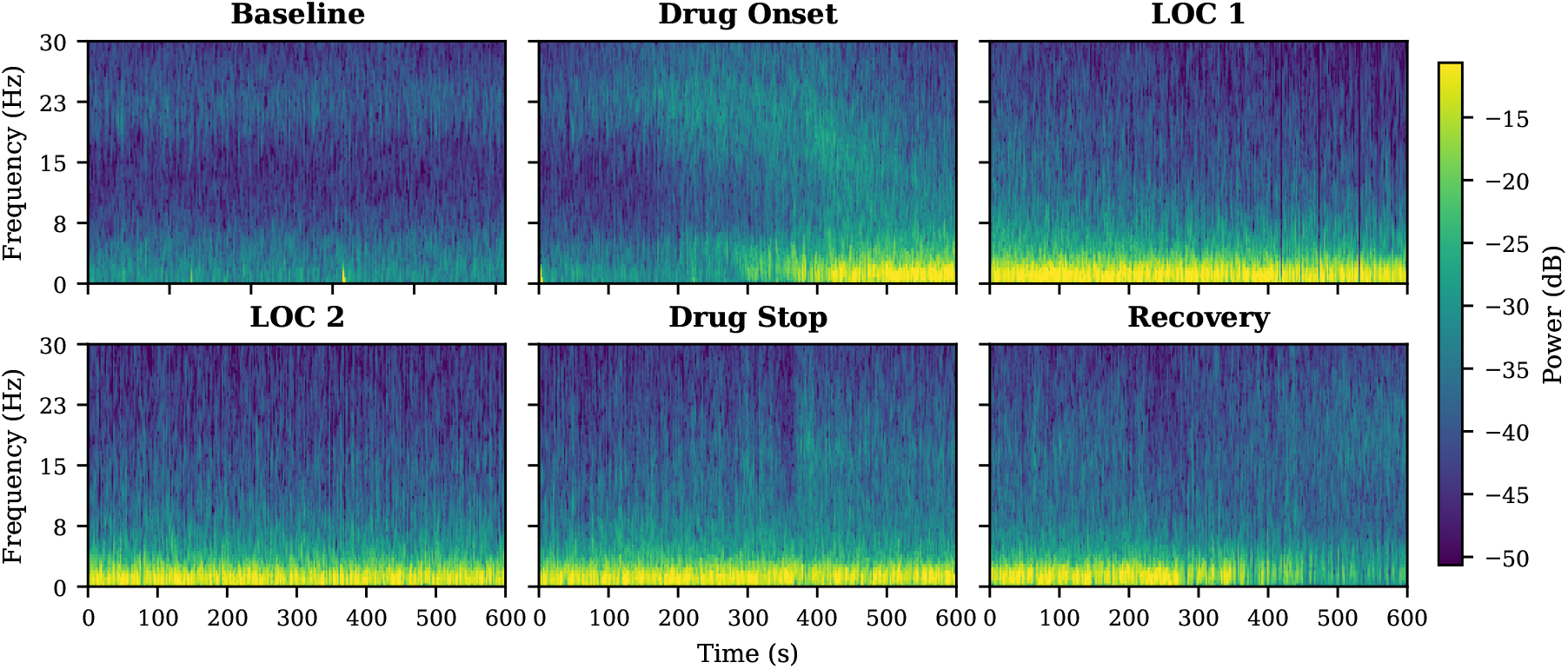
Multitaper spectrograms across anesthesia epochs. Time-frequency representations of LFP power during Baseline, Drug Onset, LOC 1, LOC 2, Drug Stop, and Recovery periods. Note the pronounced increase in low-frequency power during loss of consciousness (LOC 1–2).

**Figure S2:**
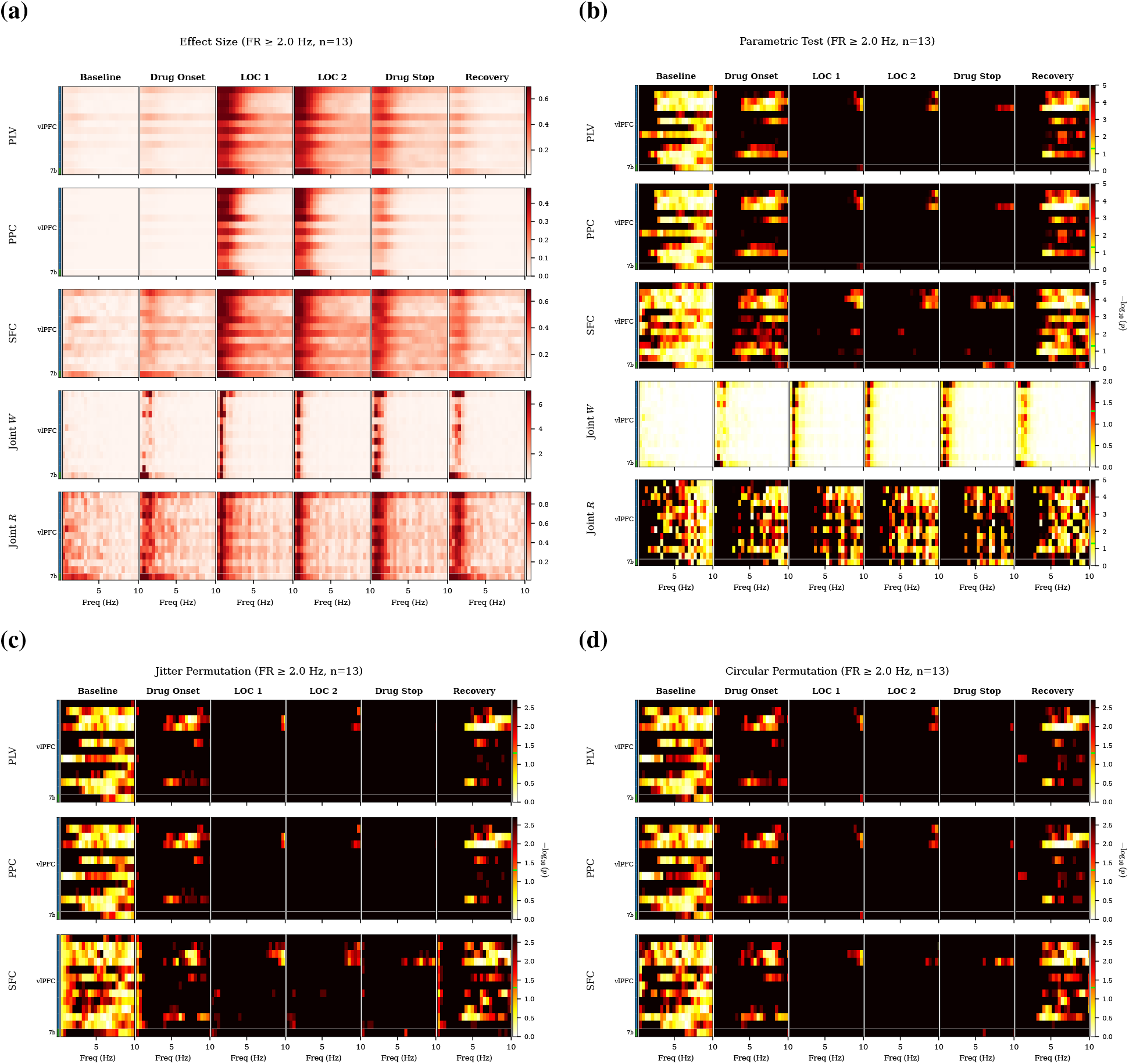
Spike–field coupling for high firing rate units (FR ≥ 2.0 Hz, *n* = 13). **(a)** Effect sizes (PLV, PPC, SFC, Joint Wald, Joint *R*) across anesthesia epochs. **(b)** Parametric *p*-values. **(c)** *p*-values from spike train jittering. (**d**) *p*-values from circular shift surrogate testing. All *p*-value heatmaps display − log_10_(*p*).

### G.2 Association learning task (a)

**Figure S3:**
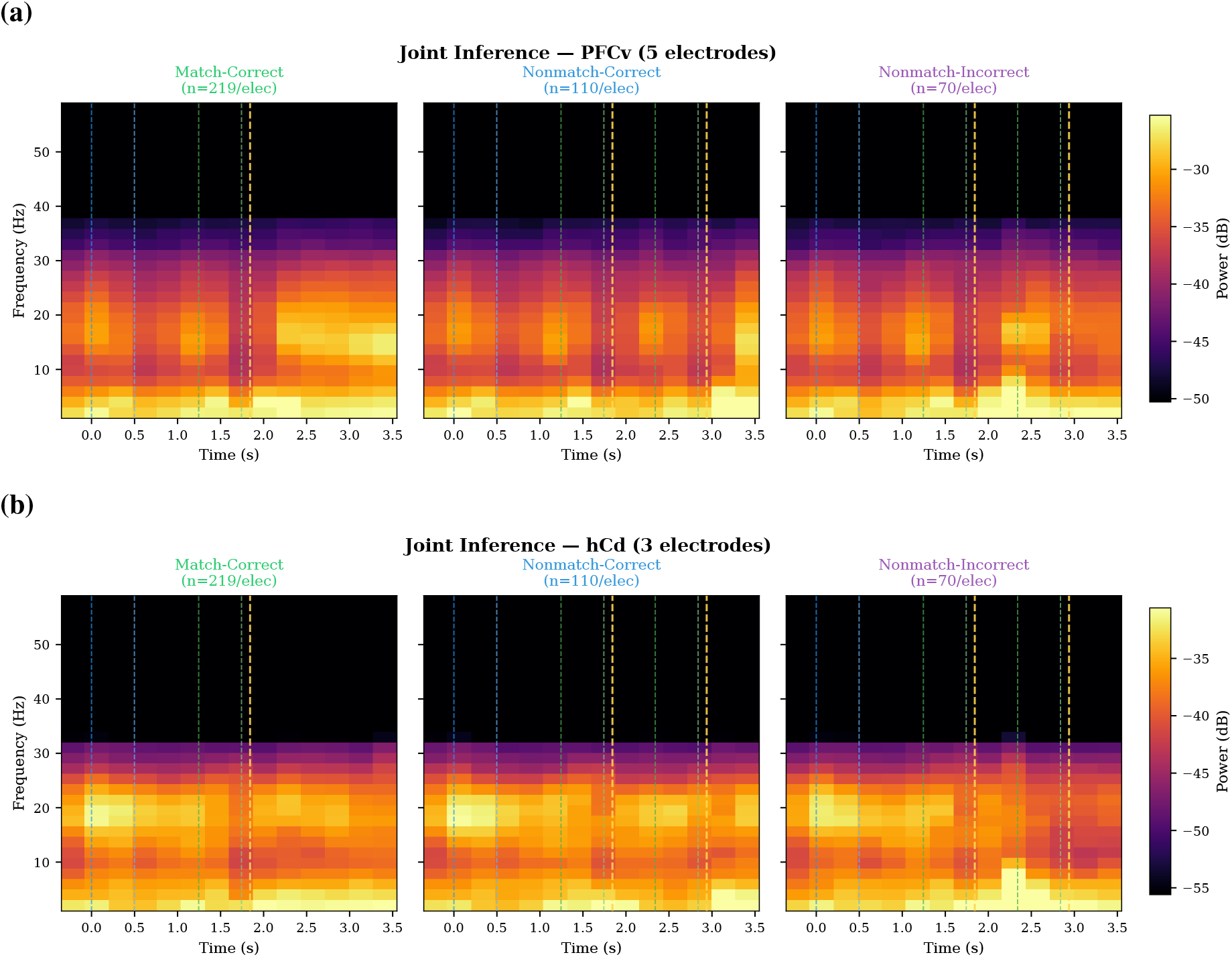
Joint inference spectrograms by brain area. Trial-averaged power spectrograms from joint spike–field inference for Match-Correct, Nonmatch-Correct, and Nonmatch-Incorrect conditions. **(a)** Area PFCv (5 electrodes). **(b)** Area hCd (3 electrodes). Dashed lines indicate task epoch boundaries.

**Figure S4:**
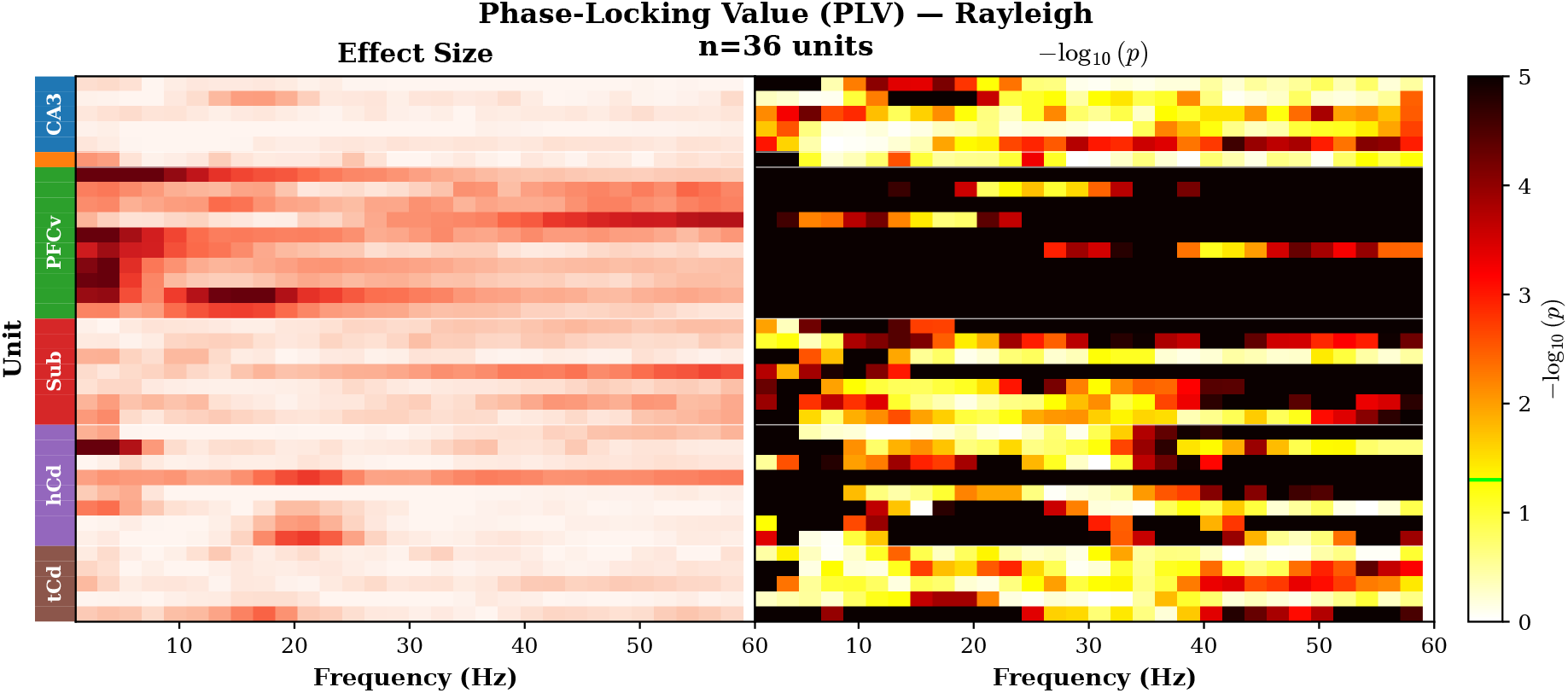
Parametric significance tests for PLV. Effect sizes (left) and Rayleigh test *p*-values (right; −log_10_ scale), sorted by brain area.

**Figure S5:**
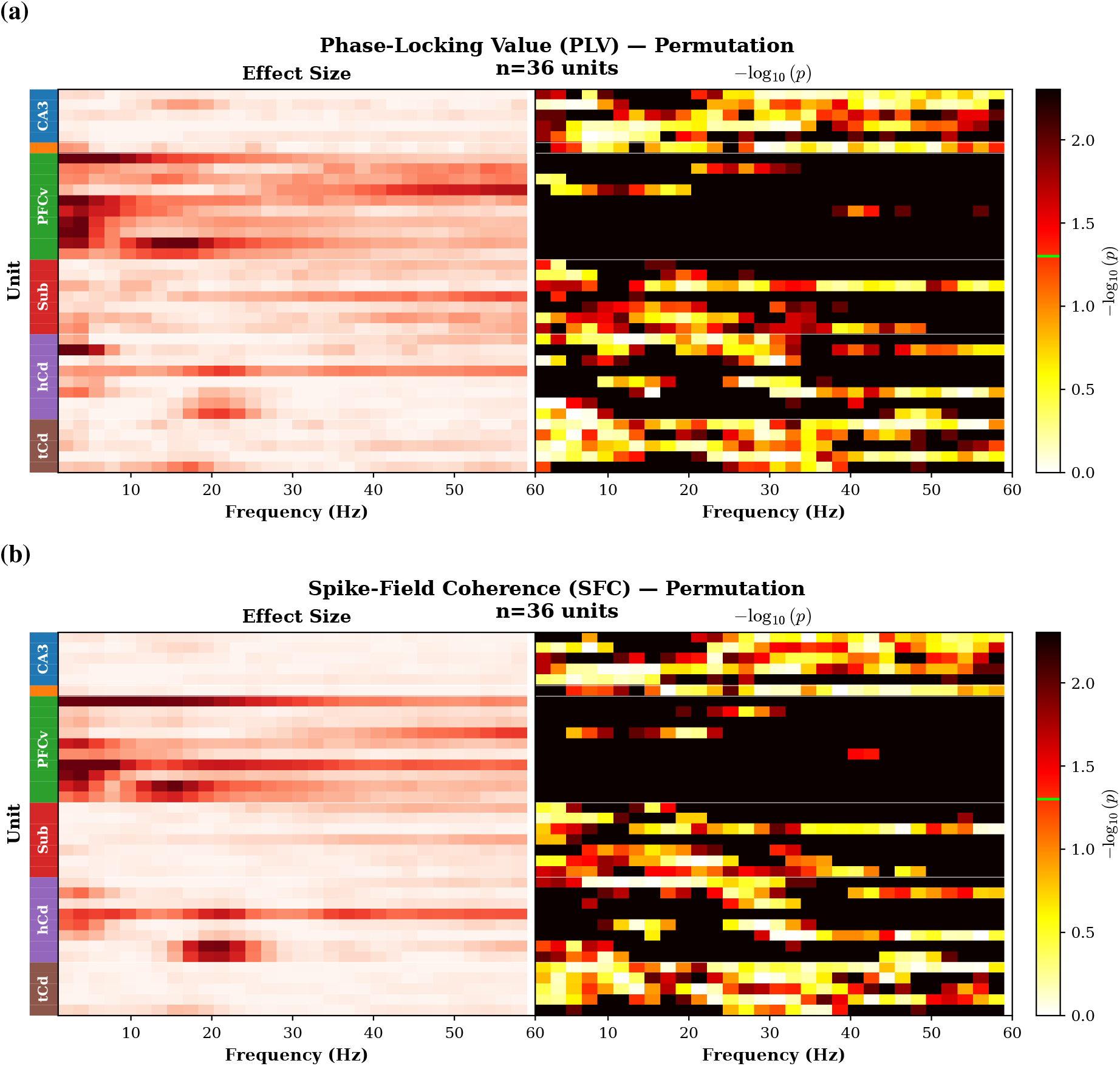
Permutation-based significance tests for spike–field coupling (*n* = 36 units). Effect sizes (left) and circular permutation *p*-values (right; −log_10_ scale), sorted by brain area. **(a)** Phase-locking value (PLV). **(b)** Spike-field coherence (SFC).

## Notes

### Competing Interest Statement

The authors have declared no competing interest.

